# Herb-paths, a network and statistical model to explore health-beneficial effects of herbs and herbal constituents

**DOI:** 10.1101/2022.10.14.512228

**Authors:** Monique van der Voet, Zhiyi Xu, Lotte van Rijnberk, Huihui Wu, Wen Liang, Pengyue Sun, Mei Wang, Marjolein Wildwater, Eefje S. Poppelaars

**Author notes:** These authors contributed equally (Monique van der Voet, Zhiyi Xu).

## Abstract

Predictions on bioactivities of herbs and herbal compounds could greatly aid drug development, but require integration of available information on herb and compound effects from various databases. We present Herb-paths, an integrated network connecting information on health-beneficial effects of herbs and herbal constituents. The statistical models included in Herb-paths allow for the calculation of the association strength of herbal (constituents) with health-beneficial effects, for both known and novel effects, and give insight into the major bioactive compounds and molecular mechanisms driving the effects.

Herb-paths’ predictions were tested and validated using a case-study of *Panax notoginseng* and its derived medicinal extract of Panaxatriol saponins (PTS), used in traditional Chinese medicine to treat stroke and other cardiovascular and cerebrovascular thromboembolic (CCT) disorders.

Results showed that Herb-paths predicted known and novel associations between PTS/*Notoginseng* and CCT phenotypes and diseases, including stroke. Predicted novel associations, such as MoyaMoya disease, are promising putative leads for treatment with PTS/*Notoginseng*, to be tested in further studies.

## INTRODUCTION

Unbiased screening for bioactivities of herbs and/or their constituents can provide a basis in the search for new drug treatments; either aiming towards the identification of a new treatment for a specific condition, or a new application of an existing treatment. Nonetheless, such an approach is expensive, time-consuming, and challenging in practice. Using available information on phenotypic and disease effects of herbs and constituents is an important tool to guide the identification of new bioactivities. However, such information is currently spread amongst various databases and no network integrates information on both health-beneficial effects of plants as a whole, as well as of their individual constituents.

Here, we present Herb-paths: a network integrating bioactivity information on plants, their constituent compounds, associated genes, and metabolic pathways. Using this integrated information, Herb-paths provides a prediction about phenotypes and diseases that might benefit from treatment with the herb or constituent, as well as insight into the molecular mechanisms of those effects. The wealth of information in Herb-paths can be used to summarize and visualize complex patterns of health-beneficial effects of herbs and their constituents. Importantly, the statistical models included in Herb-paths can estimate the strength of evidence for known effects of herbs, as well as predict novel health-beneficial effects. This provides insight into the major bioactive compounds and molecular mechanisms driving those (predicted) effects. Herb-paths could therefore be used to identify putative leads for drugs, and thereby aid in the formulation of hypotheses for testing in laboratory and clinical studies.

Given the recent interest^1^ into integrating (Western) modern medicine with the herbal knowledge of traditional Chinese medicine (TCM) that has been used in China for centuries; both Western information^2–5^ as well as Chinese information^6–8^ was integrated into Herb-paths.

Additionally, since it is common in TCM to combine several herbs into a single ‘formula’ to treat specific symptoms and diseases, this information was also integrated into Herb-paths.

To test the predictions made by the statistical models, a case-study was performed with *Panax notoginseng* (NCBI:txid44586; also known as *Radix notoginseng* or *Sanqi*). *Notoginseng* is a traditional Chinese medicinal herb, widely used for the treatment of cardiovascular and cerebrovascular thromboembolic (CCT) disorders, such as stroke^9,10^. Panaxatriol saponins (PTS) contains the major bioactive compounds extracted from Notoginseng, and is currently on the Chinese market to treat ischemic stroke^11^. PTS has been shown to have anti-platelet activity^12^, can enhance cerebral perfusion^13^, and attenuate blood-brain barrier disruption^14^. We experimentally measured the compounds present in PTS and compared their effects to those of *Notoginseng* itself. The statistical models were used to: 1) confirm whether Herb-paths indeed predicted associations between *Notoginseng* – as well as PTS compounds – and CCT diseases, and identify which compounds were responsible for which diseases, 2) validate potential new CCT diseases, and 3) visualize overarching disease associations.

## METHODS AND RESULTS

### Network creation

Data was connected from seven core databases: Dr. Duke’s Phytochemical and Ethnobotanical database^2^; FooDB^3^; Plants For A Future (PFAF)^4^; CHEBI^5^; ETCM^6^; SymMap^7^; and HERB^8^. Using R^15^ in RStudio^16^, descriptive information (e.g., names of plants, compounds, phenotypes, diseases, genes, etc.) was transformed into standardized identifiers (e.g., NCBI taxonomy^17^, international chemical shift identifier [InChI]^18^ using PubChem^19^ and ChemIDPlus^20^, Human Phenotype Ontology^21^, Human Disease Ontology^22^, Ensembl^23^,E.S. Poppelaars et al. respectively) as much as possible, in order to aid in the integration of data from different sources and allow for semantic queries. This standardization was executed using joins and regular expressions combined with manual curation.

Transforming into standardized ontology identifiers enabled the implementation of ontology-based semantic mapping (using R packages ‘ontologyIndex’ and ‘ontologyPlot’^24^), in order to include descendants of diseases in the modelling of associations with herbs and compounds, and to plot ontology trees of the results.

After integration of the different data sources, Herb-paths connects information of roughly 13.500 herbs, 3.900 formulas, 160.200 compounds, 149.200 activities, 15.400 target genes, 100 pathways, 7.600 phenotypes, 5.000 diseases, and their connections (see Figure 1).

**Figure 1.**
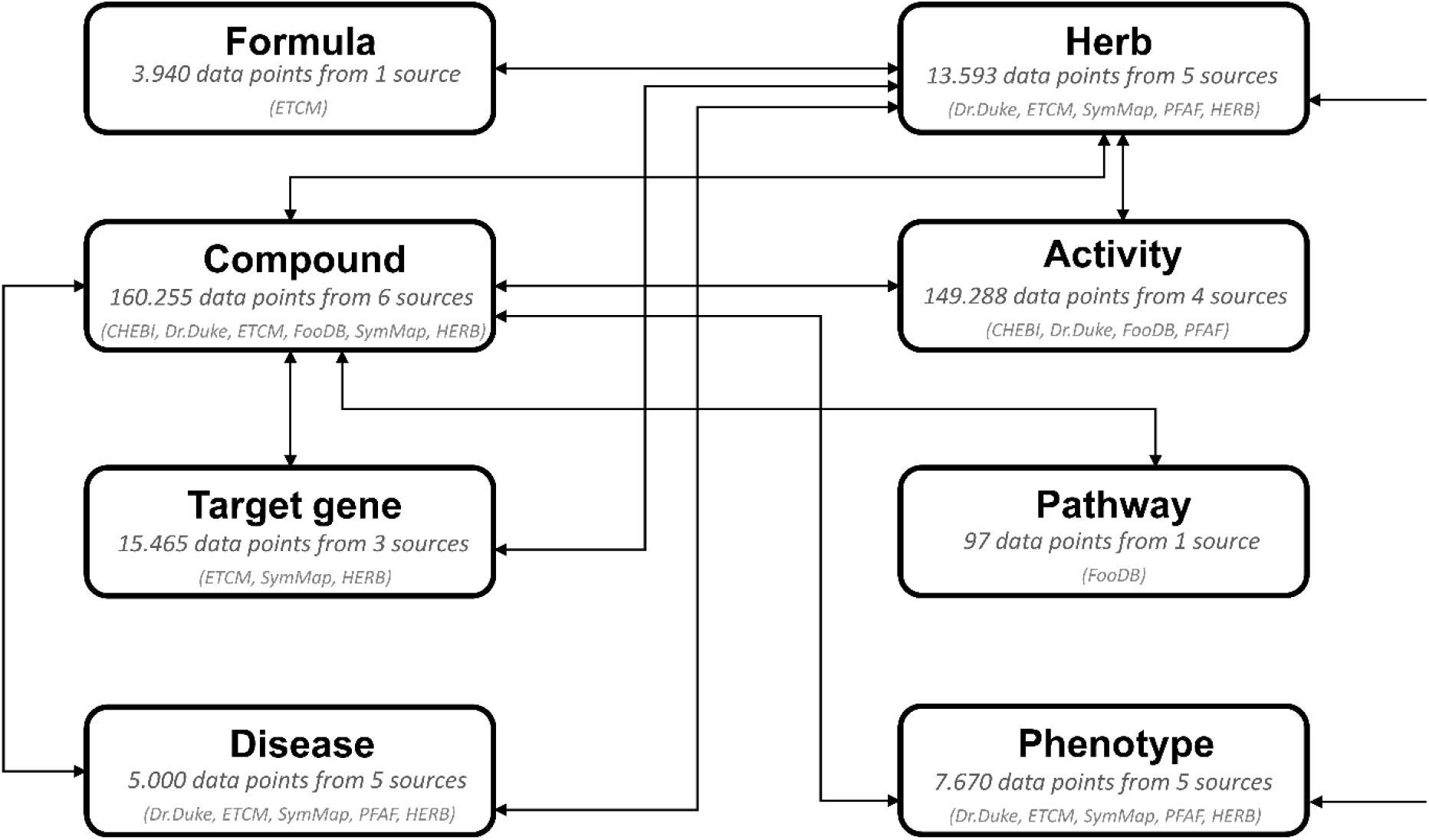
Network overview *Note*. The integrated network overview shows the eight different tables and its associations, with the number of data points and used data sources.

### *Notoginseng* and its extract Panaxatriol saponins as a case-study: Approach

To make optimal use of PTS compounds information in Herb-paths, the extracted and selected *Notoginseng* compounds present within PTS were experimentally verified using the following approaches: The candidate compounds within PTS were identified by UPLC-Q-TOF/MS (See supplementary PTS Quantitative Report), and a selection was further quantified by HPLC; including Ginsenoside Rg_1_, Ginsenoside Re, and Notoginsenoside R_1_ (see supplementary PTS Identification Report). By using these two methods, 81 compounds were identified, of which eleven compounds were analyzed with quantitative information and together were calculated to contain 88.75% of the total weight in PTS (see Table S1).

Out of the 81 unique compounds present in PTS, 49 were already included in the Herb-paths network (including ten out of eleven quantified compounds, representing at least 88.3% of the total weight in PTS) (see Table S1). Twelve of those 49 compounds were connected to *Panax notoginseng* as constituents in Herb-paths, and the remaining 37 were newly annotated. Finally, *Notoginseng* was connected to 337 compounds in total.

To assess associations with CCT disorders, terms describing CCT diseases were first standardized. Terms were manually extracted from 17 scientific articles describing the effects of PTS^11–14,25–28^ or describing CCT in general^29–37^, and transformed into identifiers of the human disease ontology^22^ (DOID) (see Figure S1 and Table S2). Ontology tree descendants of the DOID identifiers were subsequently extracted in order to include CCT diseases on all levels of specificity. The same steps were performed for CCT symptoms using the human phenotype^21^ (HP) ontology.

To examine whether *Notoginseng* constituents – and the major bioactive compounds of *Notoginseng* included in PTS – were associated with the same effects, predictions of associated diseases were modelled for both *Panax notoginseng* as a whole plant (see Figure 5A), as well as for the constituents contained within the PTS extract (see Figure 5B). Within the general disease predictions, DOID identifiers corresponding to CCT disease terms were selected. Validations of novel associated CCT diseases were performed both for all constituents of *Panax notoginseng* (see Figures S4A and 6A), as well as for the constituents contained within PTS (see Figures S4A and 6B).

### Statistical models

The Herb-paths statistical models consist of two pipelines: one for identification and another for validation (see Figure 2). Firstly, the identification pipeline estimates the association strength between herbs and diseases for known and related-yet-novel connections (see Figure 3) and returns the known associated compounds and target genes. A detailed description of the algorithm is included in the Supplementary Materials. In brief, association strength is calculated based on the disease specificity and the compound hit ratio. Disease specificity, in turn, is measured by information content score; calculated based on their ratio of total associations with compounds. Further, compound hit ratio is calculated as the number of constituents showing overlap in associations with each disease already connected to the herb, as a ratio of the total number of constituents. Next, the association strength percentage is calculated for each associated disease by multiplying the compound hit ratio with the disease specificity and calculating the percentage of the maximum association strength. Statistical significance of the association strength percentages is estimated using Monte Carlo random sampling simulations (see Figure S2), and the resulting *p*-values are corrected for multiple comparisons using the False Discovery Rate (FDR)^38^. Finally, constituents and target genes known to be associated with the predicted significant diseases are returned.

**Figure 2.**
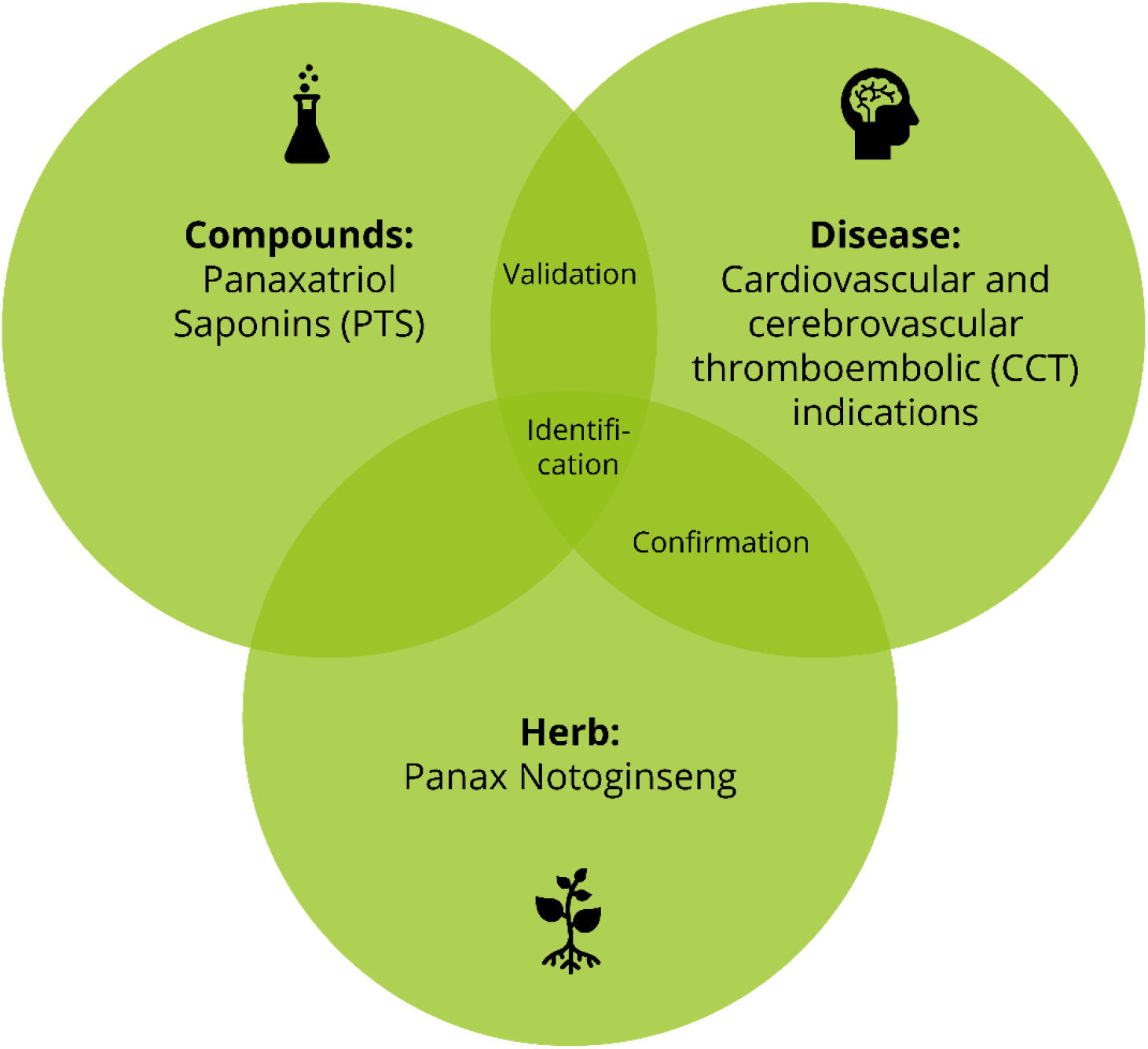
Venn diagram of statistical model approach *Note*. ‘Confirmation’ can be done at the intersection of herb and disease by summarizing existing knowledge on CCT diseases connected with Notoginseng. ‘Identification’ is performed at the intersection of herb, compound, and disease identifying responsible compounds for CCT indications by Notoginseng. ‘New CCT indications’ are explored at the intersection of compounds and diseases by using the validation pipeline. For each step, visualization of overarching associations can be performed. Phenotypes can also be modelled instead of diseases.

**Figure 3.**
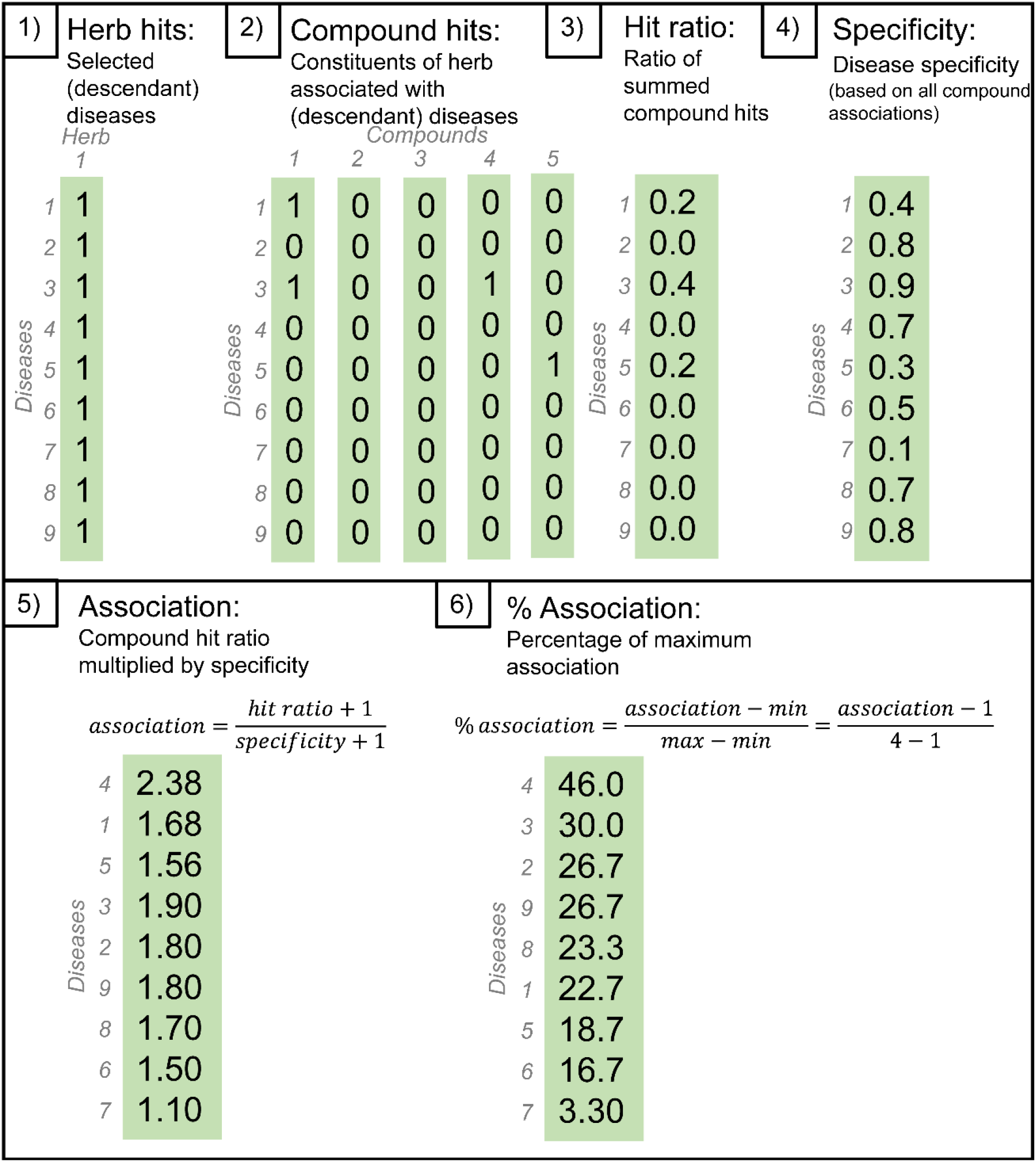
Algorithm steps of the identification pipeline to predict the association strength of diseases *Note*. The identification pipeline estimates the strength of evidence for known and novelyet-related associations between herbs and diseases, based on disease specificity (weighted information content score) and associations with the herb’s constituents. The same algorithm can be applied for phenotypes.

Secondly, the validation pipeline aims to provide validation for novel diseases associated with herb constituents (see Figure 4). A detailed description is included in the Supplementary Materials. In brief, novel diseases are identified as being associated with herb constituents but not with the herb itself. Active constituents connected with these novel diseases are retrieved. Herb similarity is calculated as the percentage of active constituent overlap. Diseases are validated by comparing the overlap between novel diseases that are associated with the input herb constituents and the diseases associated with the most similar herb.

**Figure 4.**
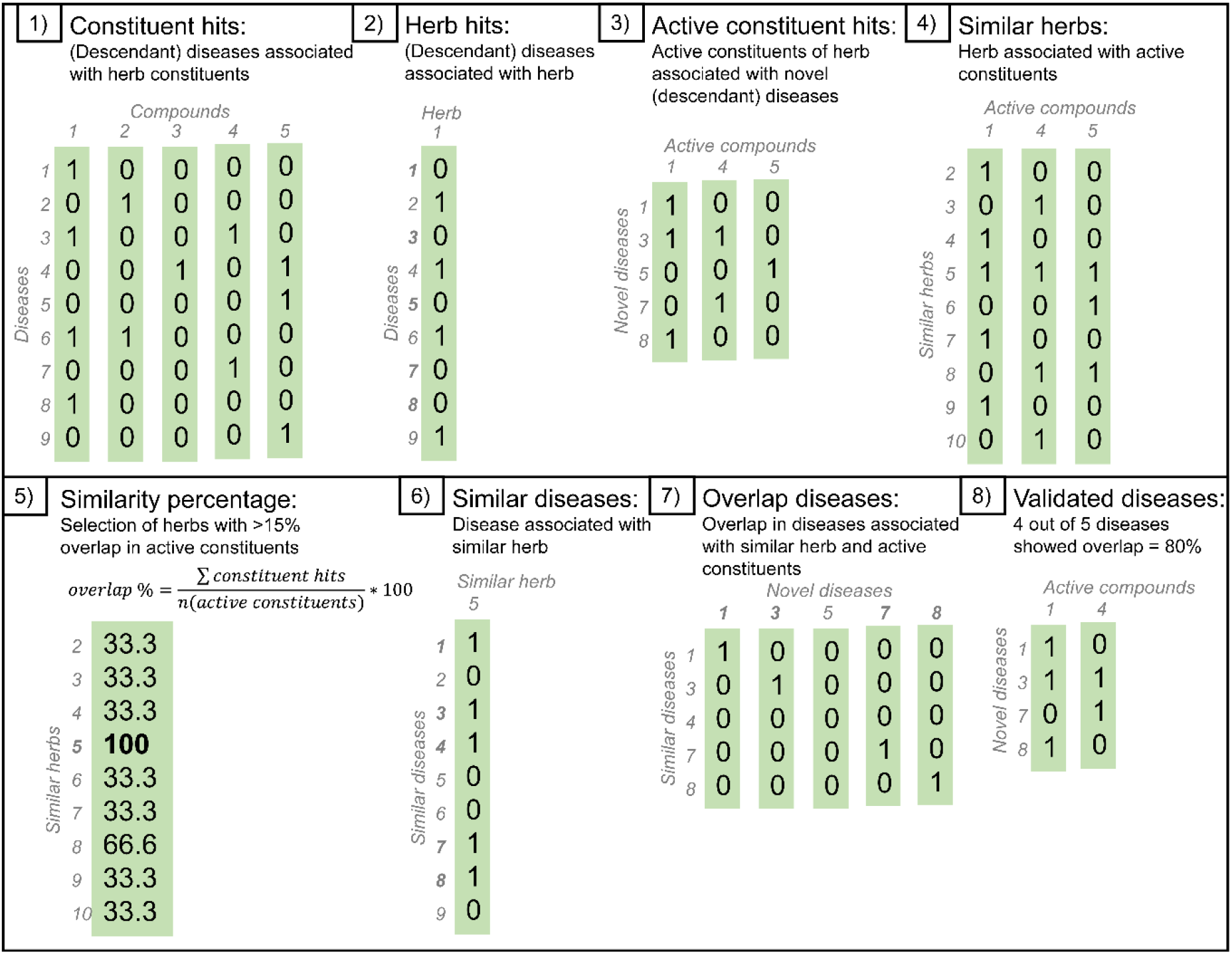
Algorithm steps of the validation pipeline to predict novel diseases *Note*. The validation pipeline predicts novel diseases associated to constituents but not the herb, and validates them based on active constituent overlap. The same algorithm can be applied for phenotypes.

Thirdly, by plotting the predicted diseases as an ontology tree with their ancestors, the complex effects of herbs can be visualized. Importantly, hub nodes can be identified that are connected with several predicted diseases (see Figures 5 and 6), thereby identifying the overarching effects of the herb as potential targets.

**Figure 5.**
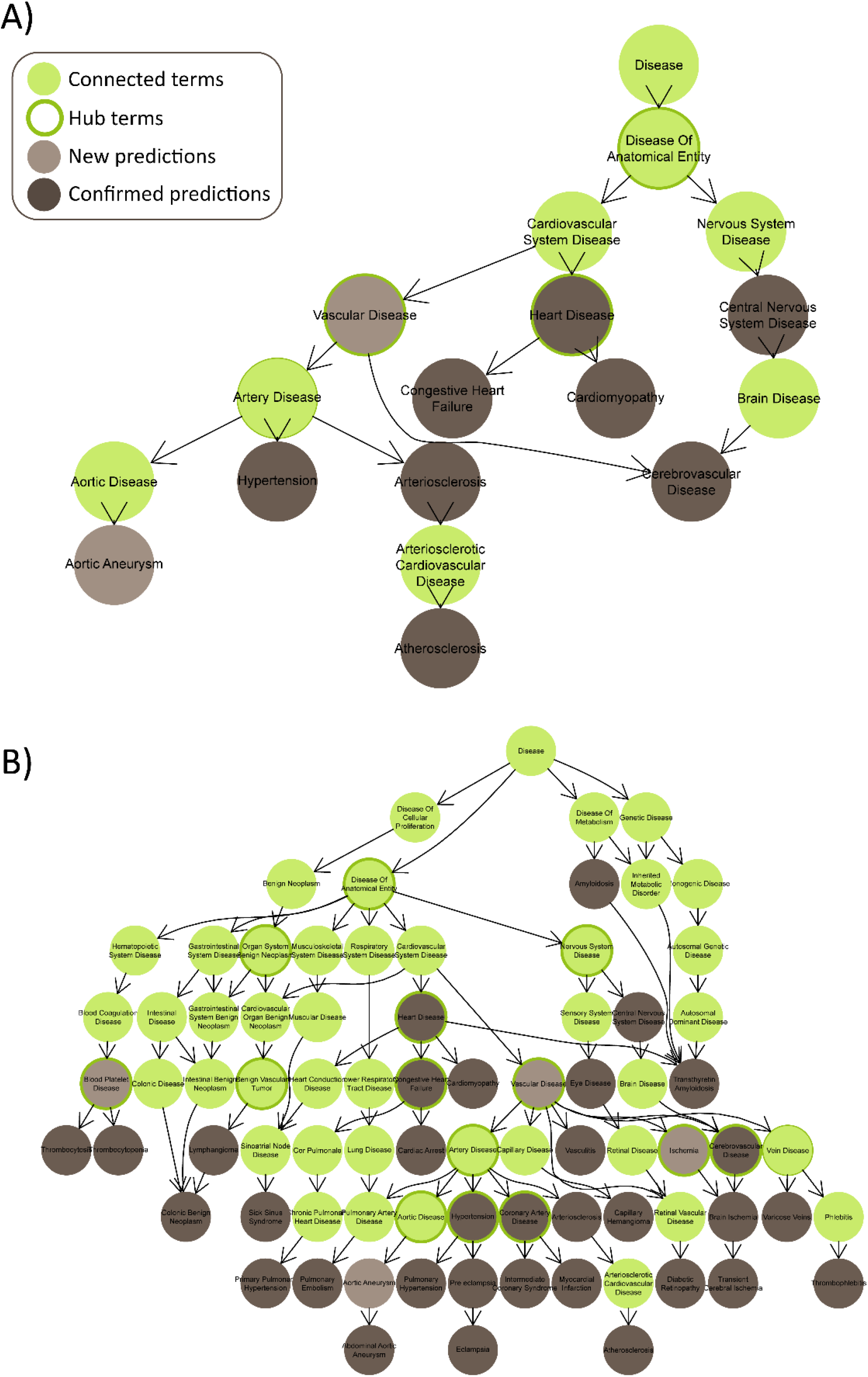
DOID ontology tree of confirmed CCT diseases *Note*. DOID ontology tree of confirmed CCT diseases using A) PTS compounds, and B) *Notoginseng*. PTS = Panaxatriol saponins; DOID = human Disease Ontology ID.

**Figure 6.**
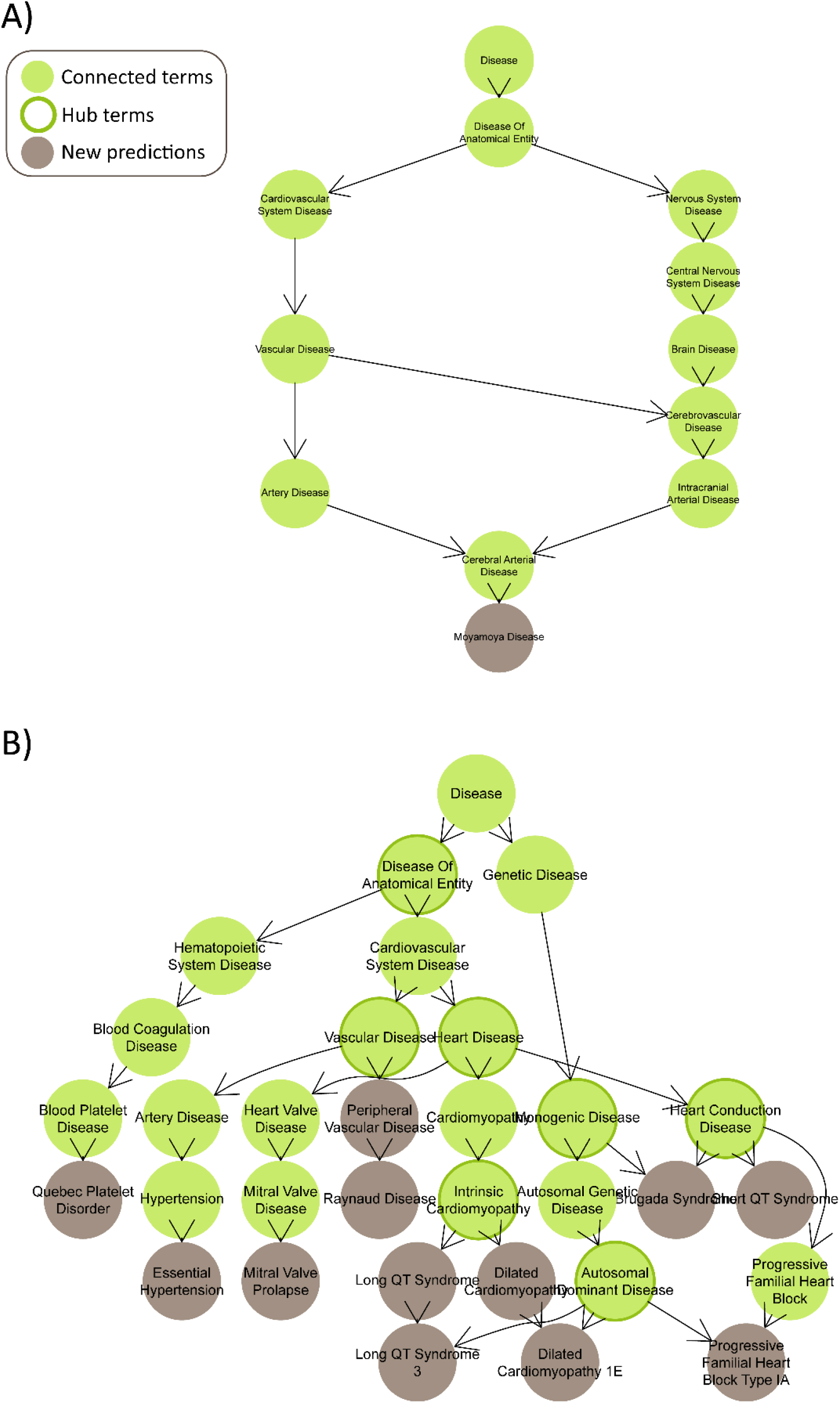
DOID ontology tree of validated new CCT diseases *Note*. DOID ontology tree of validated new CCT diseases using A) PTS compounds, and B) *Notoginseng*. PTS = Panaxatriol saponins; DOID = human Disease Ontology ID.

### Data and Software Availability

The results data is freely available via https://doi.org/10.5281/zenodo.7019208. The use of Herb-paths’ statistical models is considered proprietary and is available on request by contacting. Regardless, we believe this manuscript has unique scientific value, considering the reporting and sharing of our approach and the case-study results.

### *Notoginseng* and its extract Panaxatriol saponins as a case-study: predictions

Firstly, prediction of known and novel-yet-related associations between Panaxatriol saponins (PTS; extracted from *Notoginseng*) and CCT diseases yielded 7 known and 2 novel significant CCT diseases, with an association strength ranging from 7.4% to 12.6%. These associations included Cerebrovascular Disease and Aortic Aneurysm, and were connected via hubs of Artery Disease and Heart Disease (see Figure 5A). The associations were driven (among others) by one of the three main PTS compounds: Notoginsenoside R1.

Using all compounds connected with *Panax notoginseng*, prediction of known and novel associations between *Notoginseng* and CCT diseases yielded 30 known and 4 novel significant CCT diseases, with association strengths ranging from 7.4% to 23.4%. These associations included Cerebrovascular Disease and Ischemia, and were connected via the hubs Vascular Disease and Blood Platelet Disease (see Figure 5B). These results confirm, as expected, that both *Notoginseng* and PTS are significantly associated with CCT diseases. Additionally, while PTS is associated with several CCT diseases, *Notoginseng* itself yields stronger associations to a broader range of CCT diseases.

Secondly, validation of novel associations between compounds within PTS and CCT diseases validated one out of five novel diseases by comparing them to five herbs (see Figure S5) that all had an equal 75% overlap in active constituents and were associated with the same CCT disease: MoyaMoya disease (see Figures S4A and 6A). This association was mainly driven by three compounds: Panaxosides B and D, and Ginsenoside F4, which are all synonyms of Ginsenoside Rg1 – one of the main ingredients of PTS. Twelve target genes are involved in this association. This result demonstrates the added value of our approach for extending the indications of herbal medicine. In particular, compared to *Notoginseng*, PTS has a stronger association to Moyamoya disease.

Using all compounds connected with *Notoginseng*, validation of novel associations between constituents of *Notoginseng* (but not the herb itself) and CCT diseases validated 12 out of 26 new diseases. This was done by comparing to one herb that showed a 51% overlap in active constituents (*Tetradium ruticarpum*, see Figure S8) and was associated with the same 12 CCT diseases, such as Raynaud Disease and Quebec Platelet Disorder (see Figures S4A and 6B). The disease hubs included Vascular Disease and Heart Disease. The associations were mainly driven by 9 compounds, including the strong antioxidant Quercetin^39^. These results indicate that there are several unexplored CCT diseases that have not been previously connected with *Notoginseng* itself, but have been connected to its constituents, and could potentially be treated by *Notoginseng* and PTS. This could be an interesting starting point for further research.

Additional results for the prediction and validation of phenotypes can be found in the Supplementary Materials.

## DISCUSSION

In this study, we presented the exploration and prediction of known and novel effects of herbs and its constituents with Herb-paths: a network integrating information on health-beneficial effects and mechanisms of herbs and constituents, as well as statistical models for identification and validation of disease associations. The identification pipeline can predict the strength of associations between herbs and diseases, including known and novel-yet-related associations. Moreover, by using the validation pipeline, novel associations between herb and diseases can be explored through associations formerly only made with the herb’s constituents. The use of Herb-paths’ statistical models is available on request by contacting.

Herb-paths was tested in a case-study by using *Panax notoginseng* and the major extracted bioactive components from Panaxatriol saponins (PTS), due to its associations with cardiovascular and cerebrovascular thromboembolic (CCT) disorders^9,10^. Using the identification pipeline, 7 known CCT diseases showed significant associations to PTS, and 30 known CCT diseases showed significant associations to *Notoginseng*. These associations included the most important application of PTS: Cerebrovascular disease, specifically Brain Ischemia^9,10^; demonstrating that Herb-paths contains accurate known associations. Since the identification pipeline uses ontology-based semantic mapping, novel-yet-related diseases can also be predicted; defined as ontology tree descendants of known associations. Indeed, two novel CCT diseases showed a significant association to PTS, and four novel CCT diseases showed a significant association to *Notoginseng*. These included Aortic Aneurysm (a dilation in the aorta which can be caused/aggravated by platelet aggregation^40^); which might benefit from the anti-platelet treatment provided by *Notoginseng*/PTS^12,27^.

By plotting the predicted diseases in an ontology tree, overarching hub nodes can be identified. In all analyses, hub diseases nodes contained Vascular Disease. Similarly, Vascular Disease is indeed included in the hub nodes of all collected CCT literature terms.

Using the validation pipeline, novel associations between *Notoginseng* or PTS and diseases were explored further by using associations connected to constituents (and not the herb itself). These novel associations were validated by comparing them to effects of similar herbs (i.e., herbs with the most overlap in active constituents), that were found to have some of the same effects on diseases. Using this approach, one novel CCT disease was connected to PTS (MoyaMoya disease), while 12 diseases (including Raynaud Disease and Quebec Platelet Disorder) were connected to *Notoginseng*. Next, we will discuss a selection of the predictions in light of available scientific literature.

A novel association for treatment with PTS was MoyaMoya disease; a genetic disorder where cerebral blood flow is blocked by constrictions and thrombosis, causing ischemia and hemorrhage^41^. Although no studies have been published connecting *Notoginseng* to MoyaMoya disease, *Notoginseng* is known to treat thrombosis^42^ and have antiplatelet effects^12^ and thus could prove an effective treatment for MoyaMoya disease^41^, which should be tested in further research.

Quebec Platelet Disorder is a genetic disorder characterized by urokinase-type plasminogen activator (uPA) in platelets, which causes accelerated fibrinolysis and bleeding^43^. *Notoginseng* has been shown to have anti-platelet activities^12,27^, but an investigation in the treatment of Quebec Platelet Disorder specifically has not been published. However, one *Notoginseng* constituent (Notoginsenoside R1) has been shown to increase uPA and fibrinolysis^44^, which would aggravate Quebec Platelet Disorder; making *Notoginseng*/PTS a contraindication for Quebec Platelet Disorder. This illustrates that when associations are found by Herb-paths, it should be considered whether it is a positive or a negative association.

Finally, Raynaud Disease was predicted as a novel association to *Notoginseng*, which is a disease that results from vasospasm^45^. Vasospasm occurs when a spasm in an artery leads to vasoconstriction. Although no studies have been published connecting *Notoginseng* to vasospasm, vasospasm itself was also predicted as a novel phenotype association to *Notoginseng* (see Supplementary Materials). Importantly, both cerebral vasospasm and Raynaud disease are triggered by the same mechanism: the rho/ROCK signaling pathway^46,47^, which is inhibited by PTS^48–51^ as well as indirectly by the *Notoginseng* constituent Quercetin^52,53^. Thus, Raynaud Disease could be studied as a potential treatment target of *Notoginseng*/PTS.

In conclusion, Herb-paths is an integrated network and collection of statistical models that can summarize, explore, visualize, and predict the complex health-beneficial effects of herbs and their herbal constituents. The results of Herb-paths were validated through the case-study of *Notoginseng* and PTS.

## Supporting information

Supplementary PTS Quantitative Report

Supplementary PTS Identification Report

SMILES

## ASSOCIATED CONTENT

### Supporting Information

The following files are available free of charge:

Supplementary Materials (pdf document)

Supplementary PTS Quantitative Report (pdf document)

Supplementary PTS Identification Report (pdf document)

SMILES (xlsx file)

## AUTHOR INFORMATION

### Author Contributions (CRediT)

Conceptualization: MvdV, M.Wang, M.Wildwater; Methodology: MvdV, ZX, LvR, M.Wildwater, ESP; Software: ESP; Validation: ZX, ESP; Formal analysis: ZX, ESP; Investigation: ESP; Resources: ZX; Data Curation: MvdV, ESP; Writing – Original Draft: ESP; Writing – Review & Editing: MvdV, ZX, LvR, HW, WL, PS, M.Wang, M.Wildwater, ESP; Visualization: MvdV, ESP; Supervision: ESP; Project administration: M.Wang, M.Wildwater; Funding acquisition: MvdV, ZX, HW, WL, PS, M.Wang, M.Wildwater.

## ACKNOWLEDGMENTS

We thank the creators and contributors of our data sources: PFAF, FooDB, Dr Duke, CHEBI, SymMap, ETCM, and HERB.

## ABBREVIATIONS

CCT: Cardiovascular and cerebrovascular thromboembolic
CHEBI: Chemical Entities of Biological Interest
DARTpaths: developmental and reproductive toxicity pathways
DOID: Human disease ontology
HERB: High-throughput Experiment- and Reference-guided dataBase
HP: Human phenotype
NCBI: National library of medicine
PTS: Panaxatriol saponins.

## REFERENCES

(1) Uzuner, H.; Bauer, R.; Fan, T.-P.; Guo, D.; Dias, A.; El-Nezami, H.; Efferth, T.; Williamson, E. M.; Heinrich, M.; Robinson, N.; Hylands, P. J.; Hendry, B. M.; Cheng, Y.-C.; Xu, Q. Traditional Chinese Medicine Research in the Post-Genomic Era: Good Practice, Priorities, Challenges and Opportunities. J. Ethnopharmacol. 2012, 140 (3), 458–468. https://doi.org/10.1016/j.jep.2012.02.028.

(2) U.S. Department of Agriculture, A. R. S. Dr. Duke’s Phytochemical and Ethnobotanical Databases. http://phytochem.nal.usda.gov/. https://doi.org/10.15482/USDA.ADC/1239279.

(3) FooDB. The Metabolomics Innovation Centre: Canada 2021.

(4) Fern, K. Plants for a Future: Edible and Useful Plants for a Healthier World; Permanent Publications: Hampshire, UK, 1997.

(5) Hastings, J.; Owen, G.; Dekker, A.; Ennis, M.; Kale, N.; Muthukrishnan, V.; Turner, S.; Swainston, N.; Mendes, P.; Steinbeck, C. ChEBI in 2016: Improved Services and an Expanding Collection of Metabolites. Nucleic Acids Res. 2016, 44 (D1), D1214–D1219. https://doi.org/10.1093/nar/gkv1031.

(6) Xu, H.-Y.; Zhang, Y.-Q.; Liu, Z.-M.; Chen, T.; Lv, C.-Y.; Tang, S.-H.; Zhang, X.-B.; Zhang, W.; Li, Z.-Y.; Zhou, R.-R.; Yang, H.-J.; Wang, X.-J.; Huang, L.-Q. ETCM: An Encyclopaedia of Traditional Chinese Medicine. Nucleic Acids Res. 2019, 47 (D1), D976–D982. https://doi.org/10.1093/nar/gky987.

(7) Wu, Y.; Zhang, F.; Yang, K.; Fang, S.; Bu, D.; Li, H.; Sun, L.; Hu, H.; Gao, K.; Wang, W.; Zhou, X.; Zhao, Y.; Chen, J. SymMap: An Integrative Database of Traditional Chinese Medicine Enhanced by Symptom Mapping. Nucleic Acids Res. 2019, 47 (D1), D1110–D1117. https://doi.org/10.1093/nar/gky1021.

(8) Fang, S.; Dong, L.; Liu, L.; Guo, J.; Zhao, L.; Zhang, J.; Bu, D.; Liu, X.; Huo, P.; Cao, W.; Dong, Q.; Wu, J.; Zeng, X.; Wu, Y.; Zhao, Y. HERB: A High-Throughput Experiment- and Reference-Guided Database of Traditional Chinese Medicine. Nucleic Acids Res. 2021, 49 (D1), D1197–D1206. https://doi.org/10.1093/nar/gkaa1063.

(9) Ng, T. B. Pharmacological Activity of Sanchi Ginseng (Panax Notoginseng). J. Pharm. Pharmacol. 2010, 58 (8), 1007–1019. https://doi.org/10.1211/jpp.58.8.0001.

(10) Yang, F.; Ma, Q.; Matsabisa, M. G.; Chabalala, H.; Braga, F. C.; Tang, M. Panax Notoginseng for Cerebral Ischemia: A Systematic Review. Am. J. Chin. Med. 2020, 48 (06), 1331–1351. https://doi.org/10.1142/S0192415X20500652.

(11) He, L.; Chen, X.; Zhou, M.; Zhang, D.; Yang, J.; Yang, M.; Zhou, D. Radix/Rhizoma Notoginseng Extract (Sanchitongtshu) for Ischemic Stroke: A Randomized Controlled Study. Phytomedicine 2011, 18 (6), 437–442. https://doi.org/10.1016/j.phymed.2010.10.004.

(12) Qi, H.; Huang, Y.; Yang, Y.; Dou, G.; Wan, F.; Zhang, W.; Yang, H.; Wang, L.; Wu, C.; Li, L. Anti-Platelet Activity of Panaxatriol Saponins Is Mediated by Suppression of Intracellular Calcium Mobilization and ERK2/P38 Activation. BMC Complement. Altern. Med. 2016, 16 (1), 174. https://doi.org/10.1186/s12906-016-1160-7.

(13) Hui, Z.; Sha, D. J.; Wang, S. L.; Li, C. S.; Qian, J.; Wang, J. Q.; Zhao, Y.; Zhang, J. H.; Cheng, H. Y.; Yang, H.; Yu, L. J.; Xu, Y. Panaxatriol Saponins Promotes Angiogenesis and Enhances Cerebral Perfusion after Ischemic Stroke in Rats. BMC Complement. Altern. Med. 2017, 17 (1), 1–10. https://doi.org/10.1186/s12906-017-1579-5.

(14) Li, C.; Yang, H.; Hui, Z.; Yu, L.; Wang, S.; Chen, Y.; Xu, Y. Panaxatriol Saponins Attenuate Blood-Brain Barrier Disruption in Rats Following Transient Middle Cerebral Artery Occlusion. Int. J. Clin. Exp. Med. 2017, 10 (5), 7521–7532.

(15) R Core Team. R: A Language and Environment for Statistical Computing. R Foundation for Statistical Computing: Vienna, Austria 2022.

(16) RStudio Team Inc. RStudio: Integrated Development for R. RStudio: Boston, MA 2021.

(17) Schoch, C. L.; Ciufo, S.; Domrachev, M.; Hotton, C. L.; Kannan, S.; Khovanskaya, R.; Leipe, D.; Mcveigh, R.; O’Neill, K.; Robbertse, B.; Sharma, S.; Soussov, V.; Sullivan, J. P.; Sun, L.; Turner, S.; Karsch-Mizrachi, I. NCBI Taxonomy: A Comprehensive Update on Curation, Resources and Tools. Database 2020, 2020 (2), 1–21. https://doi.org/10.1093/database/baaa062.

(18) Heller, S. R.; McNaught, A.; Pletnev, I.; Stein, S.; Tchekhovskoi, D. InChI, the IUPAC International Chemical Identifier. J. Cheminform. 2015, 7 (1), 23. https://doi.org/10.1186/s13321-015-0068-4.

(19) Kim, S.; Chen, J.; Cheng, T.; Gindulyte, A.; He, J.; He, S.; Li, Q.; Shoemaker, B. A.; Thiessen, P. A.; Yu, B.; Zaslavsky, L.; Zhang, J.; Bolton, E. E. PubChem in 2021: New Data Content and Improved Web Interfaces. Nucleic Acids Res. 2021, 49 (D1), D1388– D1395. https://doi.org/10.1093/nar/gkaa971.

(20) Tomasulo, P. ChemIDplus-Super Source for Chemical and Drug Information. Med. Ref. Serv. Q. 2002, 21 (1), 53–59. https://doi.org/10.1300/J115v21n01_04.

(21) Köhler, S.; Gargano, M.; Matentzoglu, N.; Carmody, L. C.; Lewis-Smith, D.; Vasilevsky, N. A.; Danis, D.; Balagura, G.; Baynam, G.; Brower, A. M.; Callahan, T. J.; Chute, C. G.; Est, J. L.; Galer, P. D.; Ganesan, S.; Griese, M.; Haimel, M.; Pazmandi, J.; Hanauer, M.; Harris, N. L.; Hartnett, M. J.; Hastreiter, M.; Hauck, F.; He, Y.; Jeske, T.; Kearney, H.; Kindle, G.; Klein, C.; Knoflach, K.; Krause, R.; Lagorce, D.; McMurry, J. A.; Miller, J. A.; Munoz-Torres, M. C.; Peters, R. L.; Rapp, C. K.; Rath, A. M.; Rind, S. A.; Rosenberg, A. Z.; Segal, M. M.; Seidel, M. G.; Smedley, D.; Talmy, T.; Thomas, Y.; Wiafe, S. A.; Xian, J.; Yüksel, Z.; Helbig, I.; Mungall, C. J.; Haendel, M. A.; Robinson, P. N. The Human Phenotype Ontology in 2021. Nucleic Acids Res. 2021, 49 (D1), D1207–D1217. https://doi.org/10.1093/nar/gkaa1043.

(22) Schriml, L. M.; Mitraka, E.; Munro, J.; Tauber, B.; Schor, M.; Nickle, L.; Felix, V.; Jeng, L.; Bearer, C.; Lichenstein, R.; Bisordi, K.; Campion, N.; Hyman, B.; Kurland, D.; Oates, C. P.; Kibbey, S.; Sreekumar, P.; Le, C.; Giglio, M.; Greene, C. Human Disease Ontology 2018 Update: Classification, Content and Workflow Expansion. Nucleic Acids Res. 2019, 47 (D1), D955–D962. https://doi.org/10.1093/nar/gky1032.

(23) Howe, K. L.; Achuthan, P.; Allen, J.; Allen, J.; Alvarez-Jarreta, J.; Amode, M. R.; Armean, I. M.; Azov, A. G.; Bennett, R.; Bhai, J.; Billis, K.; Boddu, S.; Charkhchi, M.; Cummins, C.; Da Rin Fioretto, L.; Davidson, C.; Dodiya, K.; El Houdaigui, B.; Fatima, R.; Gall, A.; Garcia Giron, C.; Grego, T.; Guijarro-Clarke, C.; Haggerty, L.; Hemrom, A.; Hourlier, T.; Izuogu, O. G.; Juettemann, T.; Kaikala, V.; Kay, M.; Lavidas, I.; Le, T.; Lemos, D.; Gonzalez Martinez, J.; Marugán, J. C.; Maurel, T.; McMahon, A. C.; Mohanan, S.; Moore, B.; Muffato, M.; Oheh, D. N.; Paraschas, D.; Parker, A.; Parton, A.; Prosovetskaia, I.; Sakthivel, M. P.; Salam, A. I. A.; Schmitt, B. M.; Schuilenburg, H.; Sheppard, D.; Steed, E.; Szpak, M.; Szuba, M.; Taylor, K.; Thormann, A.; Threadgold, G.; Walts, B.; Winterbottom, A.; Chakiachvili, M.; Chaubal, A.; De Silva, N.; Flint, B.; Frankish, A.; Hunt, S. E.; IIsley, G. R.; Langridge, N.; Loveland, J. E.; Martin, F. J.; Mudge, J. M.; Morales, J.; Perry, E.; Ruffier, M.; Tate, J.; Thybert, D.; Trevanion, S. J.; Cunningham, F.; Yates, A. D.; Zerbino, D. R.; Flicek, P. Ensembl 2021. Nucleic Acids Res. 2021, 49 (D1), D884–D891. https://doi.org/10.1093/nar/gkaa942.

(24) Greene, D.; Richardson, S.; Turro, E. OntologyX: A Suite of R Packages for Working with Ontological Data. Bioinformatics 2017, 33 (7), btw763. https://doi.org/10.1093/bioinformatics/btw763.

(25) Huang, Y.; Yu, J.; Wan, F.; Zhang, W.; Yang, H.; Wang, L.; Qi, H.; Wu, C. Panaxatriol Saponins Attenuated Oxygen-Glucose Deprivation Injury in PC12 Cells via Activation of PI3K/Akt and Nrf2 Signaling Pathway. Oxid. Med. Cell. Longev. 2014, 2014, 1–11. https://doi.org/10.1155/2014/978034.

(26) Chen, Y.; Zhou, D.; Yan, S.; Liu, J.; Tan, H.; Song, Y. Study on the Mechanism of Sanqi Tongshu Capsule Combined with Butylphthalide in the Treatment of Cerebral Infarction. World Chinese Med. 2021, 16 (7). https://doi.org/10.3969/j.issn.

(27) Xu, Z.; Xu, Y.; Xie, X.; Tian, Y.; Sui, J.; Sun, Y.; Lin, D.; Gao, X.; Peng, C.; Fan, Y. Anti-Platelet Aggregation of Panax Notoginseng Triol Saponins by Regulating GP1BA for Ischemic Stroke Therapy. Chin. Med. 2021, 16 (1), 12. https://doi.org/10.1186/s13020-021-00424-3.

(28) Zhang, C.; Li, C.; Chen, S.; Li, Z.; Ma, L.; Jia, X.; Wang, K.; Bao, J.; Liang, Y.; Chen, M.; Li, P.; Su, H.; Lee, S. M. Y.; Liu, K.; Wan, J.-B.; He, C. Hormetic Effect of Panaxatriol Saponins Confers Neuroprotection in PC12 Cells and Zebrafish through PI3K/AKT/MTOR and AMPK/SIRT1/FOXO3 Pathways. Sci. Rep. 2017, 7 (1), 41082. https://doi.org/10.1038/srep41082.

(29) Flora, G. D.; Nayak, M. K. A Brief Review of Cardiovascular Diseases, Associated Risk Factors and Current Treatment Regimes. Curr. Pharm. Des. 2019, 25 (38), 4063–4084. https://doi.org/10.2174/1381612825666190925163827.

(30) Dentali, F.; Ageno, W.; Rancan, E.; Donati, A. V.; Galli, L.; Squizzato, A.; Venco, A.; Mannucci, P. M.; Manfredini, R. Seasonal and Monthly Variability in the Incidence of Venous Thromboembolism: A Systematic Review and a Meta-Analysis of the Literature. Thromb. Haemost. 2011, 106 (3), 439–447. https://doi.org/10.1160/TH11-02-0116.

(31) Laslett, L. J.; Alagona, P.; Clark, B. A.; Drozda, J. P.; Saldivar, F.; Wilson, S. R.; Poe, C.; Hart, M. The Worldwide Environment of Cardiovascular Disease: Prevalence, Diagnosis, Therapy, and Policy Issues. J. Am. Coll. Cardiol. 2012, 60 (25), S1–S49. https://doi.org/10.1016/j.jacc.2012.11.002.

(32) Townsend, N.; Wilson, L.; Bhatnagar, P.; Wickramasinghe, K.; Rayner, M.; Nichols, M. Cardiovascular Disease in Europe: Epidemiological Update 2016. Eur. Heart J. 2016, 37 (42), 3232–3245. https://doi.org/10.1093/eurheartj/ehw334.

(33) Portegies, M. L. P.; Koudstaal, P. J.; Ikram, M. A. Cerebrovascular Disease. In Handbook of Clinical Neurology; Elsevier B.V., 2016; Vol. 138, pp 239–261. https://doi.org/10.1016/B978-0-12-802973-2.00014-8.

(34) Medrano Albero, M. J.; Boix Martínez, R.; Cerrato Crespán, E.; Ramírez Santa-Pau, M. Incidence and Prevalence of Ischaemic Heart Disease and Cerebrovascular Disease in Spain: A Systematic Review of the Literature. Rev. Esp. Salud Publica 2006, 80 (1), 05– 15. https://doi.org/10.1590/S1135-57272006000100002.

(35) Tong, X.; Yang, Q.; Ritchey, M. D.; George, M. G.; Jackson, S. L.; Gillespie, C.; Merritt, R. K. The Burden of Cerebrovascular Disease in the United States. Prev. Chronic Dis.2019, 16 (4), 1–9. https://doi.org/10.5888/pcd16.180411.

(36) Ghanima, W.; Brodin, E.; Schultze, A.; Shepherd, L.; Lambrelli, D.; Ulvestad, M.; Ramagopalan, S.; Halvorsen, S. Incidence and Prevalence of Venous Thromboembolism in Norway 2010–2017. Thromb. Res. 2020, 195 (June), 165–168. https://doi.org/10.1016/j.thromres.2020.07.011.

(37) Deitelzweig, S. B.; Johnson, B. H.; Lin, J.; Schulman, K. L. Prevalence of Clinical Venous Thromboembolism in the USA: Current Trends and Future Projections. Am. J. Hematol. 2011, 86 (2), 217–220. https://doi.org/10.1002/ajh.21917.

(38) Benjamini, Y.; Yekutieli, D. The Control of the False Discovery Rate in Multiple Testing under Dependency. Ann. Stat. 2001, 29 (4), 1165–1188. https://doi.org/10.1214/aos/1013699998.

(39) Shaik, Y. B.; Castellani, M. L.; Perrella, A.; Conti, F.; Salini, V.; Tete, S.; Madhappan, B.; Vecchiet, J.; De Lutiis, M. A.; Caraffa, A.; Cerulli, G. Role of Quercetin (a Natural Herbal Compound) in Allergy and Inflammation. J. Biol. Regul. Homeost. Agents 2006, 20 (3–4), 47–52.

(40) Sun, W.; Zheng, J.; Gao, Y. Targeting Platelet Activation in Abdominal Aortic Aneurysm: Current Knowledge and Perspectives. Biomolecules 2022, 12 (2), 206. https://doi.org/10.3390/biom12020206.

(41) Fujimura, M.; Bang, O. Y.; Kim, J. S. Moyamoya Disease. In Frontiers of Neurology and Neuroscience; 2016; Vol. 40, pp 204–220. https://doi.org/10.1159/000448314.

(42) Zhu, C.; Jiang, H.; Zhou, X.; Zhang, Z.; Wu, Y.; Fang, H.; Wang, L. Blood Circulation Activating Effect of Sanqi (Radix Notoginseng) on Venous Thromboembolism Rat. J. Tradit. Chinese Med. 2021, 41 (5), 753–761. https://doi.org/10.19852/j.cnki.jtcm.20201023.001.

(43) Hayward, P. M.; Georges E Rivard, &. Quebec Platelet Disorder. Expert Rev. Hematol. 2011, 4 (2), 137–141. https://doi.org/10.1586/ehm.11.5.

(44) Zhang, W. J.; Wojta, J.; Binder, B. R. Effect of Notoginsenoside R1 on the Synthesis of Components of the Fibrinolytic System in Cultured Smooth Muscle Cells of Human Pulmonary Artery. Cell. Mol. Biol. (Noisy-le-grand). 1997, 43 (4), 581–587.

(45) Temprano, K. K. A Review of Raynaud’s Disease. Mo. Med. 2016, 113 (2), 123–126.

(46) Sato, M.; Tani, E.; Fujikawa, H.; Kaibuchi, K. Involvement of Rho-Kinase–Mediated Phosphorylation of Myosin Light Chain in Enhancement of Cerebral Vasospasm. Circ. Res. 2000, 87 (3), 195–200. https://doi.org/10.1161/01.RES.87.3.195.

(47) Fardoun, M. M.; Nassif, J.; Issa, K.; Baydoun, E.; Eid, A. H. Raynaud’s Phenomenon: A Brief Review of the Underlying Mechanisms. Front. Pharmacol. 2016, 7 (NOV), 1– 13. https://doi.org/10.3389/fphar.2016.00438.

(48) Shi, X.; Yu, W.; Yang, T.; Liu, W.; Zhao, Y.; Sun, Y.; Chai, L.; Gao, Y.; Dong, B.; Zhu, L. Panax Notoginseng Saponins Provide Neuroprotection by Regulating NgR1/RhoA/ROCK2 Pathway Expression, in Vitro and in Vivo. J. Ethnopharmacol. 2016, 190, 301–312. https://doi.org/10.1016/j.jep.2016.06.017.

(49) Yang, T.; Miao, Y.; Zhang, T.; Mu, N.; Ruan, L.; Duan, J.; Zhu, Y.; Zhang, R. Ginsenoside Rb1 Inhibits Autophagy through Regulation of Rho/ROCK and PI3K/MTOR Pathways in a Pressure-Overload Heart Failure Rat Model. J. Pharm. Pharmacol. 2018, 70 (6), 830–838. https://doi.org/10.1111/jphp.12900.

(50) He, K.; Yan, L.; Pan, C.-S.; Liu, Y.-Y.; Cui, Y.-C.; Hu, B.-H.; Chang, X.; Li, Q.; Sun, K.; Mao, X.-W.; Fan, J.-Y.; Han, J.-Y. ROCK-Dependent ATP5D Modulation Contributes to the Protection of Notoginsenoside NR1 against Ischemia-Reperfusion-Induced Myocardial Injury. Am. J. Physiol. Circ. Physiol. 2014, 307 (12), H1764– H1776. https://doi.org/10.1152/ajpheart.00259.2014.

(51) Zheng, H.; Fu, X.; Shang, J.; Lu, R.; Ou, Y.; Chen, C. Ginsenoside Rg1 Protects Rat Bone Marrow Mesenchymal Stem Cells against Ischemia Induced Apoptosis through MiR-494-3p and ROCK-1. Eur. J. Pharmacol. 2018, 822, 154–167. https://doi.org/10.1016/j.ejphar.2018.01.001.

(52) Tao, X.; Chen, Q.; Li, N.; Xiang, H.; Pan, Y.; Qu, Y.; Shang, D.; Go, V. L. W.; Xue, J.; Sun, Y.; Zhang, Z.; Guo, J.; Xiao, G. G. Serotonin-RhoA/ROCK Axis Promotes Acinar-to-Ductal Metaplasia in Caerulein-Induced Chronic Pancreatitis. Biomed. Pharmacother. 2020, 125 (December 2019), 109999. https://doi.org/10.1016/j.biopha.2020.109999.

(53) Lai, W.-W.; Hsu, S.-C.; Chueh, F.-S.; Chen, Y.-Y.; Yang, J.-S.; Lin, J.-P.; Lien, J.-C.; Tsai, C.-H.; Chung, J.-G. Quercetin Inhibits Migration and Invasion of SAS Human Oral Cancer Cells through Inhibition of NF-?B and Matrix Metalloproteinase-2/-9 Signaling Pathways. Anticancer Res. 2013, 33 (5), 1941–1950.

