## Supplementary Materials for "Herb-paths, a network and statistical model to explore health-beneficial effects of herbs and herbal constituents"

### *Notoginseng* and its extract Panaxatriol saponins as a case-study: Approach

**Table S1**. Compounds measured in PTS

| Compound name | CAS number | Weight (%) | In Herb-paths | Connected with Notoginseng |
|---|---|---|---|---|
| Ginsenoside Rg1 | 22427-39-0 | 57.35 | Yes | Yes |
| Notoginsenoside R1 | 80418-24-2 | 16.44 | Yes | Yes |
| Ginsenoside Re | 52286-59-6 | 8.05 | Yes | Yes |
| Ginsenoside Rb1 | 41753-43-9 | 3.02 | Yes | Yes |
| Notoginsenoside R2 | 80418-25-3 | 0.63 | Yes | No |
| R-Notoginsenoside R2 | 948046-15-9 | 0.09 | Yes | Yes |
| Ginsenoside Rh1 | 63223-86-9 | 0.63 | Yes | Yes |
| R-Ginsenoside Rh1 | 80952-71-2 | 0.33 | Yes | No |
| Ginsenoside F2 | 62025-49-4 | 1.16 | Yes | Yes |
| Ginsenoside Rk3 | 364779-15-7 | 0.41 | No | |
| Ginsenoside Rh4 | 174721-08-5 | 0.64 | Yes | No |
| L-tryptophan | 73-22-3 | | Yes | No |
| 3- Methylphenyl 6- O- β- D- xylopyranosyl- β- D- glucopyranoside | 2131787-98-7 | | No | |
| Notoginsenoside J | 193977-11-6 | | Yes | Yes |
| (3β, 6β, 12β) - 20- (β- D- Glucopyranosyloxy) - 3, 12, 24, 25- tetrahydroxydammaran - 6- yl 6- O- β- D- xylopyranosyl- β- D- glucopyranoside | 1133882-74-2 | | No | |
| Vinaginsenoside R22 | 226713-28-6 | | Yes | No |
| (3β, 6α, 12β) - 3, 12, 24, 25- Tetrahydroxy- 20- (β- D- glucopyranosyloxy) dammaran- 6- yl O- pentosyl- β- D- glucopyranoside | 1394905-99-7 | | No | |
| Isoconiferoside | 1347015-00-2 | | No | |
| (3β, 6β, 12β) - 20- (β- D- Glucopyranosyloxy) - 3, 12, 24, 25- tetrahydroxydammaran - 6- yl 6- O- (6- deoxy- α- L- mannopyranosyl) - β- D- glucopyranoside | 1133882-75-3 | | No | |
| Baimaside | 18609-17-1 | | Yes | No |
| Floralginsenoside C | 942586-98-3 | | No | |
| Notoginsenoside H | 193976-69-1 | | Yes | Yes |
| Notoginsenoside SP1 | 1801723-21-6 | | No | |
| Notoginsenoside SP2 | 1801723-22-7 | | No | |
| Quercetin 3-O-sambubioside | 83048-35-5 | | Yes | No |
| Kaempferol 3,7-diglucoside | 25615-14-9 | | Yes | No |
| Floralginsenoside B | 942586-97-2 | | No | |
| Hyperoside | 482-36-0 | | Yes | No |
| Vina- ginsenoside R25 | 340270-89-5 | | No | |
| Kaempferol 3-O-sambubioside | 27661-51-4 | | No | |
| Isoquercetin | 482-35-9 | | Yes | No |
| (3β, 6β, 12β) - 6, 20- Bis(β- D- glucopyranosyloxy) - 3, 12- dihydroxydammar- 25- en- 24- one | 1135445-37-2 | | No | |

| | | | |
|---|---|---|---|
| Eugenol rutinoside | 138772-01-7 | No | |
| Gypenoside LXIX | 109150-50-7 | Yes | No |
| Gypenoside LVI | 105214-48-0 | Yes | No |
| Notoginsenoside N | 350586-56-0 | Yes | No |
| Floranotoginsenoside A | 1179351-11-1 | No | |
| Vinaginsenoside R8 | 156042-22-7 | Yes | No |
| Gypenoside XLIII | 94705-68-7 | Yes | No |
| Notoginsenoside M | 394246-74-3 | Yes | No |
| Notoginsenoside-R6 | 87741-78-4 | Yes | Yes |
| Ginsenoside Re4 | 1255210-79-7 | No | |
| 20-Glucoginsenoside Rf | 68406-27-9 | Yes | No |
| (3β, 6β, 12β) - 20- (β- D- Glucopyranosyloxy) - 3, 12, 25- trihydroxydammaran- 6- yl 2- O- β- D- glucopyranosyl- β- D- glucopyranoside | 1133882-87-7 | No | |
| Quinquenoside L16 | 454686-08-9 | No | |
| Yesanchinoside D | 163403-91-6 | No | |
| Notoginsenoside A | 193895-21-5 | Yes | No |
| Vinaginsenoside R13 | 156398-72-0 | Yes | No |
| Pseudoginsenoside RT3 | 98474-75-0 | Yes | No |
| Ginsenoside III | 223710-06-3 | Yes | No |
| Yesanchinoside D isomer | | No | |
| Ginsenoside F5 | 189513-26-6 | Yes | No |
| Notoginsenoside Fh7 | 2172820-99-2 | No | |
| Notoginsenoside Fh3 | 2172820-95-8 | No | |
| Ginsenoside F3 | 62025-50-7 | Yes | No |
| Notoginsenoside B | 193895-26-0 | Yes | Yes |
| (6β, 12β) - 6, 20- Bis(β- D- glucopyranosyloxy) - 12- hydroxydammar- 24- en- 3- one | 1614215-08-5 | No | |
| Vina-ginsenoside R4 | 156009-80-2 | Yes | No |
| Ginsenoside Rf | 52286-58-5 | Yes | Yes |
| Notoginsenoside D | 193895-50-0 | Yes | No |
| Gypenoside L | 94987-09-4 | Yes | No |
| Chikusetsusaponin VI | 137348-15-3 | Yes | No |
| Notoginsenoside Fa | 88100-04-3 | Yes | No |
| 20(S)-Ginsenoside Rg2 | 80952-72-3 | Yes | No |
| 20(R)-Ginsenoside Rg2 | 80952-72-3 | No | |
| Malonyl-ginsenoside Rb1 | 88140-34-5 | Yes | No |
| Quinquenoside R1 | 88140-34-5 | No | |
| Ginsenoside F1 | 53963-43-2 | Yes | No |
| Chikusetsusaponin IV | 7518-22-1 | Yes | No |
| 20S- Sanchirhinoside A2 | 1702363-34-5 | No | |
| Ginsenoside Rd | 52705-93-8 | Yes | No |
| 6- acetate(3β, 6α, 12β) - 3, 12, 20- Trihydroxydammar- 24- en- 6- yl β- D- glucopyranoside | 784213-07-6 | No | |

| | | | |
|---|---|---|---|
| (3β, 6β, 12β) - 6- (Acetyloxy) - 3, 12- dihydroxydammar- 24- en- 20- yl β- D- glucopyranoside | 1133882-72-0 | No | |
| (3β, 12β) - 12- Hydroxy- 6- (D- xylopyranosyloxy) dammara- 20, 24(or 20(22) , 24) - dien- 3- yl β- D- glucopyranoside | 1375079-47-2 | No | |
| Ginsenoside Rg4 | 126223-28-7 | Yes | No |
| (3β, 12β) - 12- Hydroxy- 3- (D- xylopyranosyloxy) dammara- 20, 24(or 20(22) , 24) - dien- 6- yl β- D- glucopyranoside | 1375079-55-2 | No | |
| Ginsenoside Rg6 | 147419-93-0 | Yes | No |
| Ginsenoside Rg3 | 14197-60-5 | Yes | No |
| 20(R) - Ginsenoside Rs3 | 203849-13-2 | No | |
| Ginsenoside Rs3 | 194861-70-6 | No | |
| Ginsenoside Rg5 | 186763-78-0 | Yes | No |

*Note*. SMILES of the compounds are available via the supplement ‘SMILES.csv’.

**Figure S1.** Flowchart of steps taken to select ontology terms for CCT indications

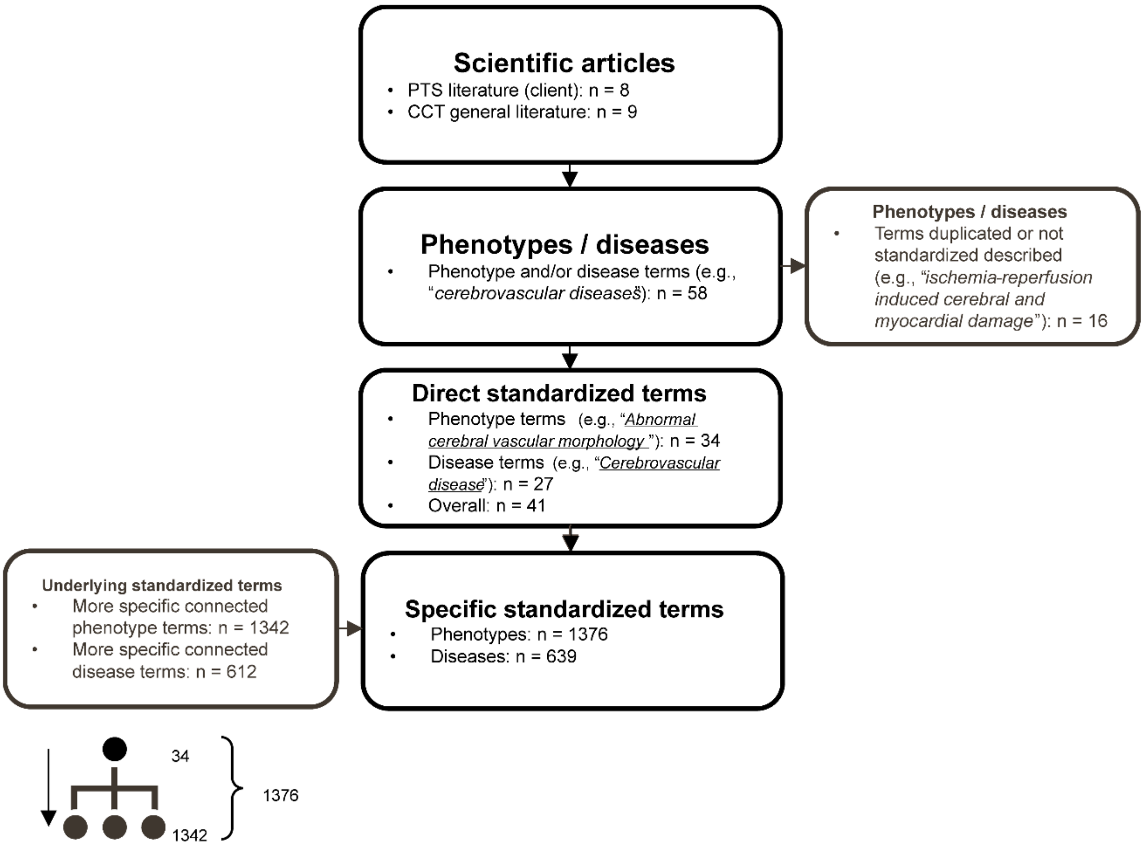

**Table S2.** Cardiovascular and cerebrovascular thromboembolic (CCT) ontology terms

| Literature CCT term | HP name | DOID name |
|---|---|---|
| ischemic stroke | Ischemic stroke | |
| stroke | Stroke | |
| ischemic stroke of anterior cerebral circulation | | anterior cerebral artery infarction |
| ischemic diseases | Tissue ischemia | ischemia |
| ischemia-reperfusion injury | Encephalomalacia | encephalomalacia |
| ischemic brain injury | Brain atrophy | |
| cerebral ischemia and reperfusion | | |
| oxygen-glucose deprivation-reperfusion induced oxidative injury | | |
| ischemia-reperfusion induced cerebral and myocardial damage | | |
| oxygen-glucose deprivation-reperfusion induced cell death | | |
| Blood-brain barrier disruption after ischemia/reperfusion | | |
| biphasic permeability of blood-brain barrier | | |
| cerebral ischemia | Cerebral ischemia | brain ischemia |
| focal cerebral ischemia | | |
| post-ischemic inflammation | | |
| cerebral edema | Cerebral edema | |
| brain edema | | brain edema |
| vasogenic edema | | |
| transient middle cerebral artery occlusion | Middle cerebral artery stroke | middle cerebral artery infarction |
| infarct expansion | | |
| cerebral infarction | | brain infarction |
| hemorrhagic transformation | | |
| cerebrovascular diseases | Abnormal cerebral vascular morphology | cerebrovascular disease |
| cerebral vascular diseases | | |
| cardiac vascular diseases | Abnormality of the cardiovascular system | cardiovascular system disease |
| vascular diseases | Abnormality of the vasculature | vascular disease |
| increased platelet aggregation | Spontaneous platelet aggregation | blood platelet disease |
| intravascular thrombosis | Venous thrombosis | thrombosis |
| vascular stenosis | Arterial stenosis | coronary stenosis |
| cerebral artery obstruction | Cerebral artery stenosis | cerebral artery occlusion |
| oxidative stress | | |
| cell apoptosis | | |
| inflammatory cytokine release | Cytokine storm | |
| acute coronary syndrome | Acute coronary syndrome | |
| 6-OHDA-induced dopaminergic neuron loss | Neuronal loss in basal ganglia | |
| coronary artery disease | Coronary artery atherosclerosis | coronary artery disease |
| coronary heart disease | | heart disease |
| venous thromboembolism | Thromboembolism | |

| | | |
|---|---|---|
| venous sinus thrombosis | Central venous sinus thrombosis | intracranial sinus thrombosis |
| deep-vein thrombosis | Deep venous thrombosis | |
| pulmonary embolism | Pulmonary embolism | pulmonary embolism |
| peripheral vascular disease | | peripheral vascular disease |
| myocardial infarction | Myocardial infarction | myocardial infarction |
| cardiac arrhythmia | Arrhythmia | |
| ischemic heart disease | | |
| hemorrhagic stroke | Cerebral hemorrhage | |
| subarachnoid hemorrhage | Subarachnoid hemorrhage | |
| arterial dissection | Arterial dissection | aortic dissection |
| Cerebral Autosomal Dominant Arteriopathy with Sub-cortical Infarcts and Leukoencephalopathy | | CADASIL |
| subcortical infarct | | |
| leukoencephalopathy | Leukoencephalopathy | |
| arteriovenous malformation | Arteriovenous malformation | arteriovenous malformation |
| moyamoya disease | Moyamoya phenomenon | Moyamoya disease |
| silent infarct | | silent myocardial infarction |
| white-matter lesion | Focal white matter lesions | |
| microbleed | | |
| transient ischemic attack | Transient ischemic attack | transient cerebral ischemia |
| primary central nervous system vasculitis | Cerebral vasculitis | central nervous system vasculitis |

*Note.* HP = Human Phenotype ontology; DOID = Human Disease Ontology.

### METHODS OF STATISTICAL MODELS

#### Specificity of diseases

The specificity of diseases is calculated using the following steps:

1) The information content (IC) score is calculated for each disease based on the number of compounds associations (see equation 1). Here, $f$ represents the number of compounds that the current disease is associated with, whereas $n$ represents the total number of associations between all diseases and all compounds (i.e., a constant). (For example, an aspecific disease such as “central nervous system disease” that occurs

4,639 times has an IC of 6.88, while a more specific disease such as "Autosomal recessive spinocerebellar ataxia 19" that occurs only once has an IC of 19.06.)

$$\mathrm{IC} = -\log_2\left(\frac{f}{n}\right) \quad (1)$$

2) A weighted IC is calculated so that it ranges from 0 (i.e., least specific) to 1 (i.e., most specific) and is thereby normalized (see equation 2). Thus, the specific IC value that would become a 0 or 1 depends on the distribution of IC values for the dataset.

$$ICweight = \frac{IC - min(IC)}{max(IC) - min\,(IC)} \quad (2)$$

3) The same calculations can be done for phenotypes instead of diseases.

**Disease association-strength estimation**

The identification pipeline uses an algorithm to find the diseases that are associated with both a herb and its constituents, and estimate its association strength (see Figure 3), before identifying responsible compounds and target genes. It does this by using the following steps:

1) A single herb is represented by a vector of all diseases that are either associated with it or not (1 = hit and 0 = no hit). Associations with descendant diseases are also counted as a hit.
2) The compounds that are contained in the herb (i.e., constituents) are looked up, and the diseases that overlap between herb and compound association are marked (1 = hit and 0 = no hit). Associations with descendant diseases are also counted as a hit. It is also possible here to make a selection of relevant constituents.
3) The compound *hit ratio* is calculated by summing the compound hits for each disease, and then calculating the ratio of summed compounds hits, compared to the total number of the herb's constituents.

4) Each disease also has a *specificity* value (i.e., weighted information score of all associated compounds for that disease, regardless of herbs; see the ‘Specificity of diseases’ section).

5) An *association strength* score is calculated by multiplying the compound *hit ratio* with the *specificity* score (see equation 3). The resulting *association strength* score ranges between 1 and 4.

$$association\ score = \frac{compound\ hit\ ratio + 1}{specificity + 1} \quad (3)$$

6) To get the *association strength percentage*, the percentage of the maximum *association strength* score is calculated, taking the theoretical minimum and maximum score (of 1 and 4) into account (see equation 4). This *association strength percentage* simplifies interpretation by magnifying differences to add nuance within a clearer range (0-100).

$$\%\ \max association\ score = \frac{score - minimum}{maximum - minimum} = \frac{score - 1}{4 - 1} \quad (4)$$

7) The same calculations can be done for phenotypes instead of diseases.

8) The associated constituents and target genes for each disease are returned as well.

**Disease association-strength significance**

*p*-values of predicted association strength of diseases are calculated by using Monte Carlo simulations (see Figure S2). It does this by calculating the chance of false positive results, using the following steps:

1) Do 1000 simulations to randomly draw the same number of compounds as there were constituents. (For example, say you have calculated the *association strength percentage* of 100 diseases connected to one specific herb with 200 constituents, then a simulation will be done a 1000 times by drawing a random set of 200 compounds

and calculating the association strength for each of the 100 diseases, given those 200 random ‘constituents’.)

2) Calculate the *association strength percentage* for each disease connected with the input herb based on overlap with the herb’s ‘constituents’. This is done both for the true set of constituents as well as for the random set of constituents for each simulation.

3) Calculate the number of false positive associations by counting the number of simulations where the random *association strength percentage* was equal to or larger than the actual *association strength percentage*, i.e. the disease was equally or more likely to be associated at random.

4) Calculate the *p*-value as the chance of false positives, by using a ratio between the percentage of false positive associations over the number of simulations (equation 5). For example, say that after 1000 simulations, 50 simulations showed an equal or larger *association strength percentage* than the actual *association strength percentage*, i.e., there are 50 positives, then the *p*-value becomes 0.051 (i.e., 50+1/1000+1).

$$p = \frac{\text{Number of false positives+1}}{\text{Total number of simulations+1}} \quad (5)$$

5) The *p*-values for all diseases are then corrected for multiple comparisons using the false discovery rate (Benjamini & Hochberg, 1995). FDR-corrected *p*-values < .05 are considered significant.

6) The same calculations can be done for phenotypes instead of diseases.

**Figure S2**. Algorithm steps of the identification pipeline to estimate significance of predictions

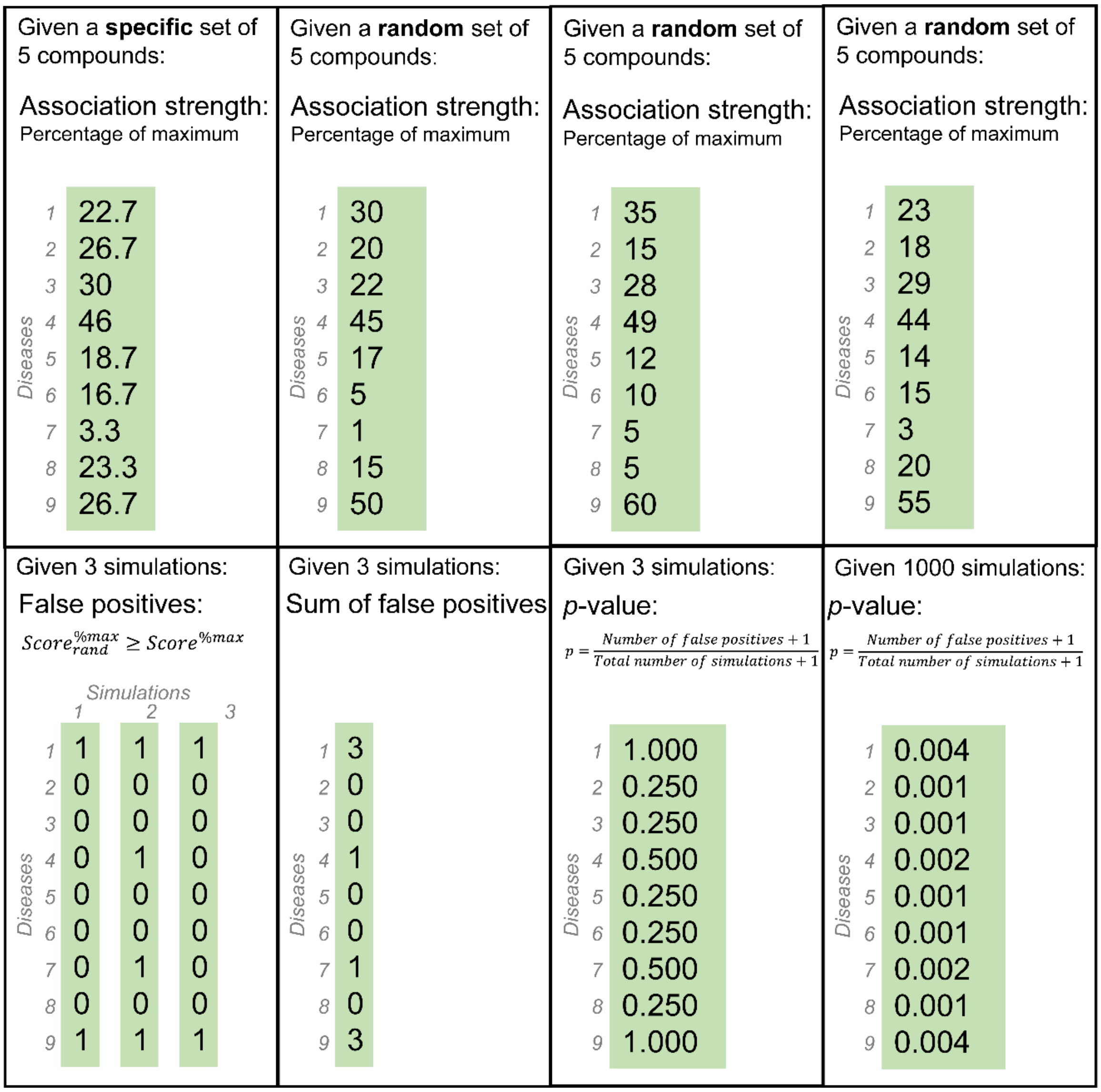


*Note.* Estimates of significance were performed through 100 simulations of Monte Carlo random sampling.

### Validation of novel CCT diseases

The validation pipeline uses an algorithm to find novel diseases that are associated with the herb constituents but not the herb itself, and validates them by finding the overlap with diseases associated with a similar herb (i.e., with the same active constituents), before retrieving responsible compounds and target genes (see Figure 4). It does this by using the following steps (see also Figure S4):

1) Look up the constituents of a specific herb, and find the diseases associated with those compounds. Associations with descendant diseases are also taken into account. It is also possible here to make a selection of relevant diseases.
2) Look up the diseases associated with the input herb.
3) Remove all diseases that are also associated with the input herb. Remove all constituents not responsible for the novel diseases to retain only active constituents.
4) The herbs that contain (some of) the same constituents are looked up, and the number of compounds that overlap between similar herbs and active constituents are counted (1 = overlap and 0 = no overlap).
5) The percentage of overlap relative to the total number of active constituents is calculated. Of the herbs that contain at least a 15% overlap in active constituents, the herb(s) with the highest number of overlap is selected.
6) The diseases that are associated with the similar herb(s) are looked up, and the diseases that do not overlap with the novel diseases connected with the active constituents are removed.
7) The diseases that overlap between those associated with the similar herb(s) and those associated with the active constituents are counted (1 = overlap and 0 = no overlap). The percentage of overlap relative to the total number of novel diseases connected with the active constituents is calculated. Each disease that showed overlap in associations to the similar herb(s) is considered validated.
8) The associated constituents and target genes for each validated disease are returned as well.
9) The same calculations can be done for phenotypes instead of diseases.

### Additional figures

The NCBI Taxonomy ontology tree for input herbs and similar herbs (based on overlap in constituents) were plotted for each run of the validation pipeline (see Figures S5, S7, and S8). These plots show the herb similarity based on plant family.

### *Notoginseng* and its extract Panaxatriol saponins as a case-study: Predictions

Firstly, prediction of known and novel-yet-related associations between PTS and phenotypes yielded 12 known and 9 novel significant cardiovascular and cerebrovascular thromboembolic (CCT) phenotypes, with association strength ranging from 7.2% to 13.9%. Associations included Stroke and Ventricular Arrhythmia, and hubs included Vasculature and Brain Morphology (see Figure S3A). These associations were driven (among others) by two of the three main PTS compounds (Ginsenoside Re and Ginsenoside Rg1).

Expanding the compound selection, prediction of known and novel associations between *Panax notoginseng* and CCT phenotypes yielded 62 known and 19 novel-yet-related significant phenotypes; with association strength ranging from 7.2% to 20.1%. These associations included Stroke and Transient Ischemic Attack, and hubs included Vasculature and Arrhythmia (see Figure S3B).

Secondly, validation of novel associations between constituents of *Notoginseng* (but not the herb itself) and CCT phenotypes validated six out of 15 new phenotypes. This was done by comparing to one herb that showed a 52% overlap in connected constituents (*Panax Ginseng*[37], see Figure S7) and was associated with the same six CCT phenotypes, such as Vasospasm and Shock (see Figure S6). The associations were mainly driven by seven compounds (see Table S3). Phenotype hubs included Vasculature and Cardiovascular System Physiology. There were no novel phenotypes that could be validated for associations with PTS compounds.

Shock comes in four major categories with different causes and treatments, but one thing they all have in common is a disturbance to blood flow[43]. Since *Notoginseng* can increase blood flow[13,42], some types of shock might benefit from *Notoginseng* treatment. Further research is necessary to explore this.

**Figure S3.** HP ontology tree of confirmed CCT phenotypes

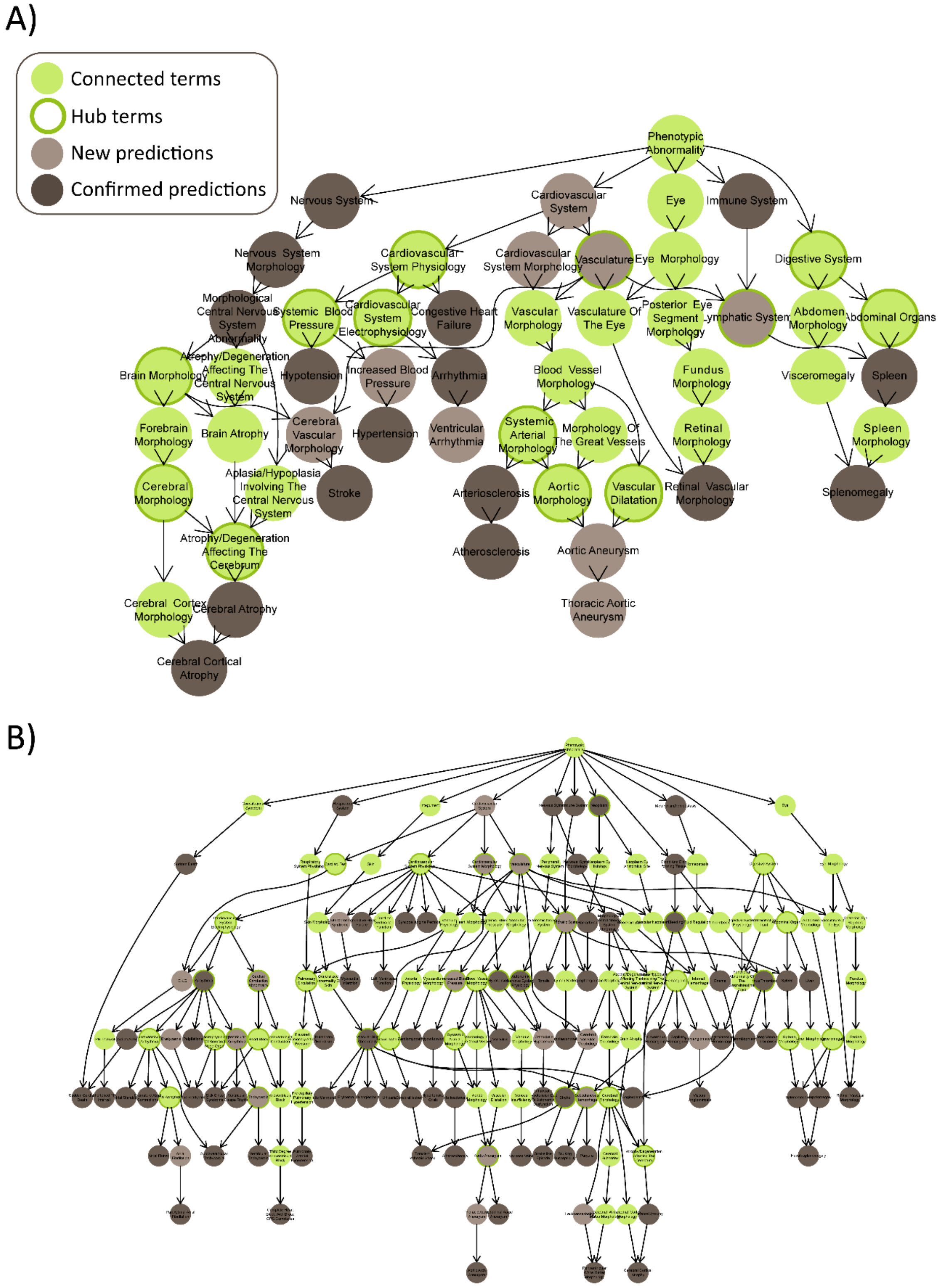

*Note*. HP ontology tree of confirmed CCT phenotypes using A) PTS compounds, and B) *Notoginseng*.

**Figure S4.** Flowchart of algorithm results of new CCT indications validation

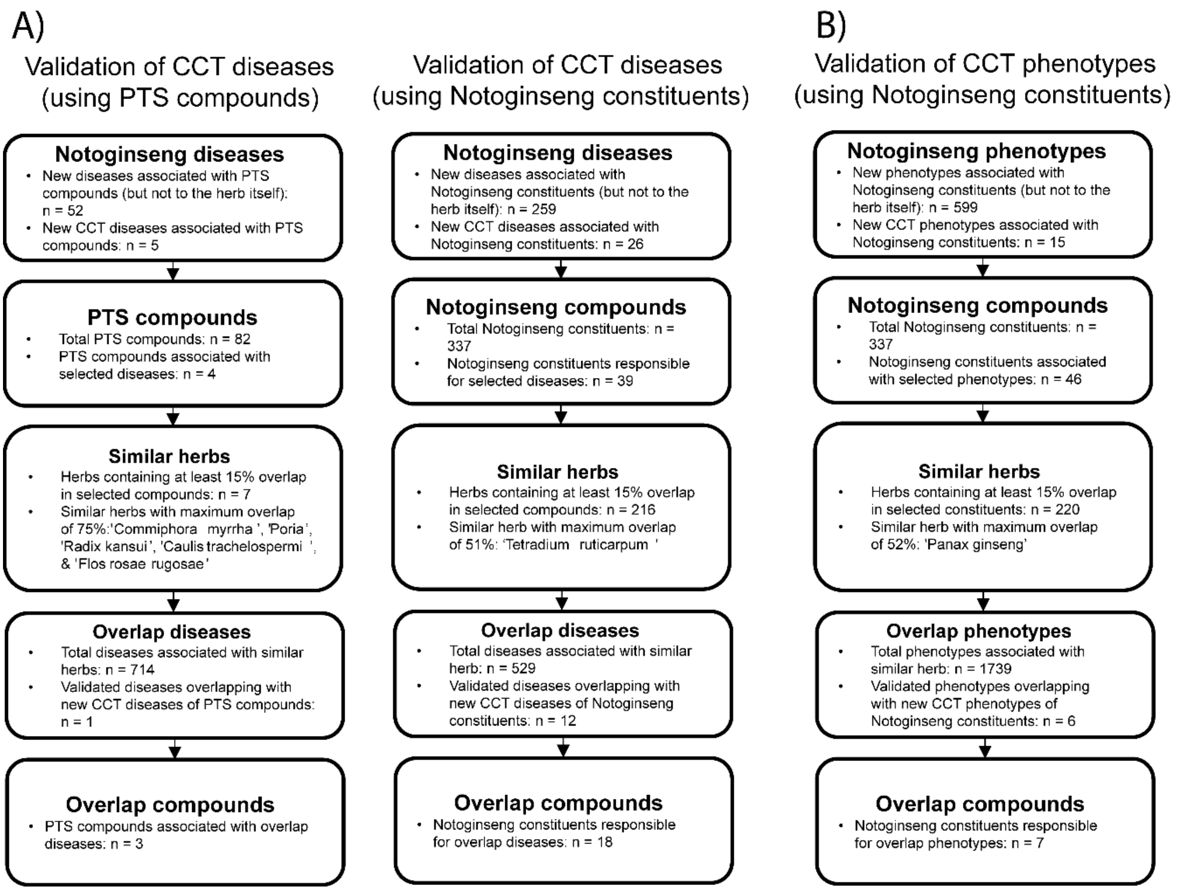


*Note*. A) Algorithm results of new CCT diseases validation; B) Algorithm results of new CCT phenotypes validation.

**Figure S5.** NCBI ontology tree of herb similarity for validation of new CCT diseases using PTS compounds

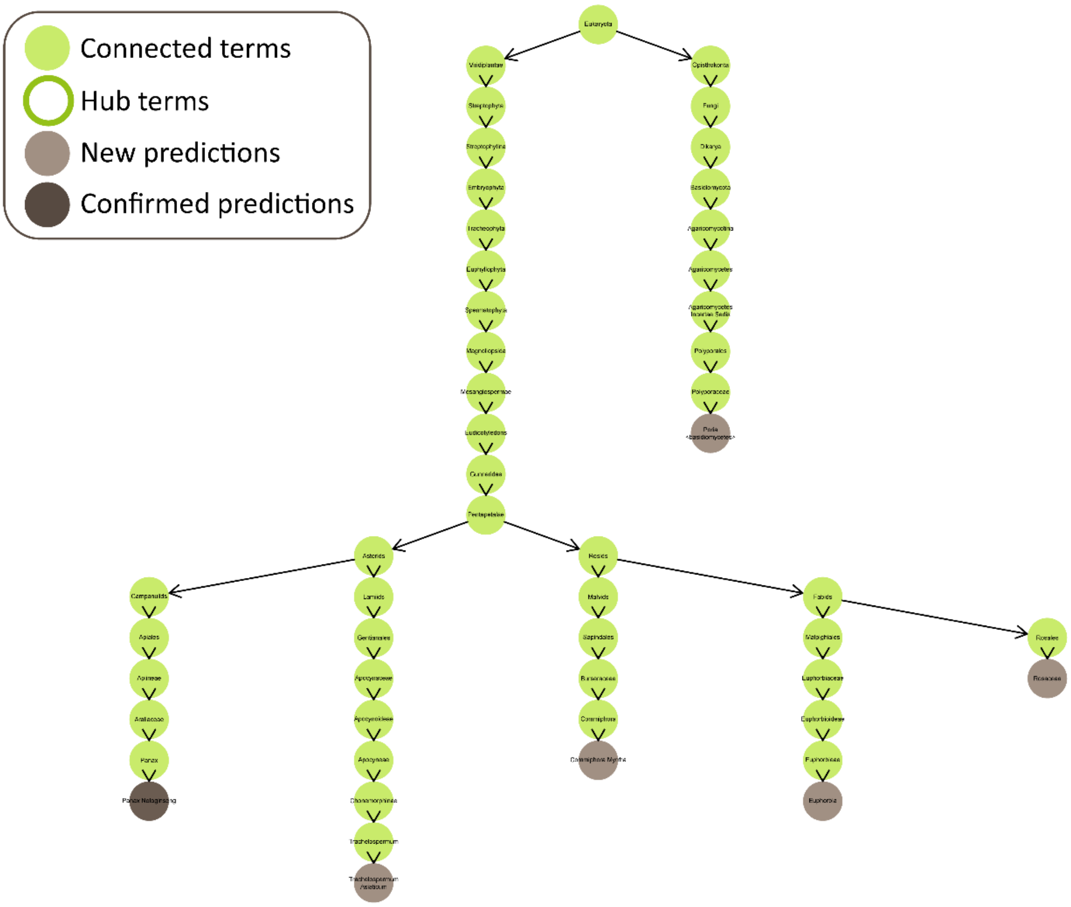

**Figure S6.** HP ontology tree of validated new CCT phenotypes using *Notoginseng*

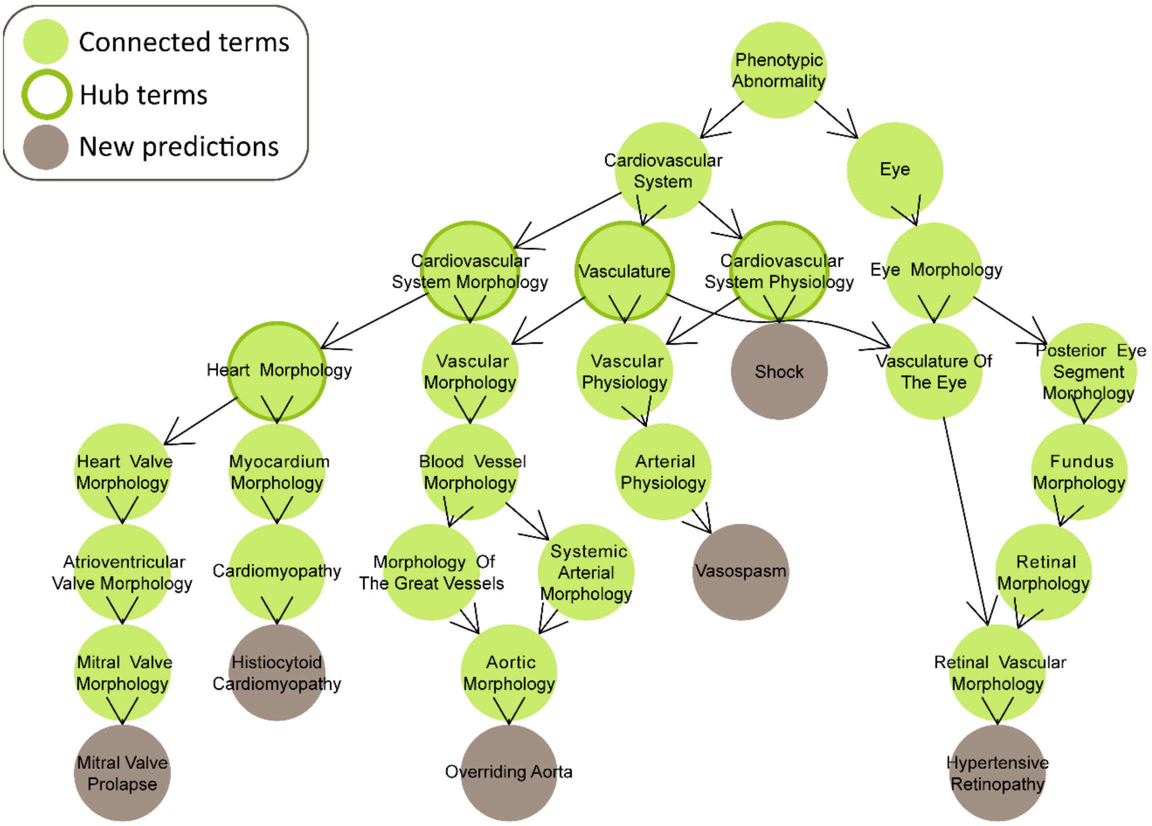

**Figure S7.** NCBI ontology tree of herb similarity for validation of new CCT phenotypes using *Notoginseng*

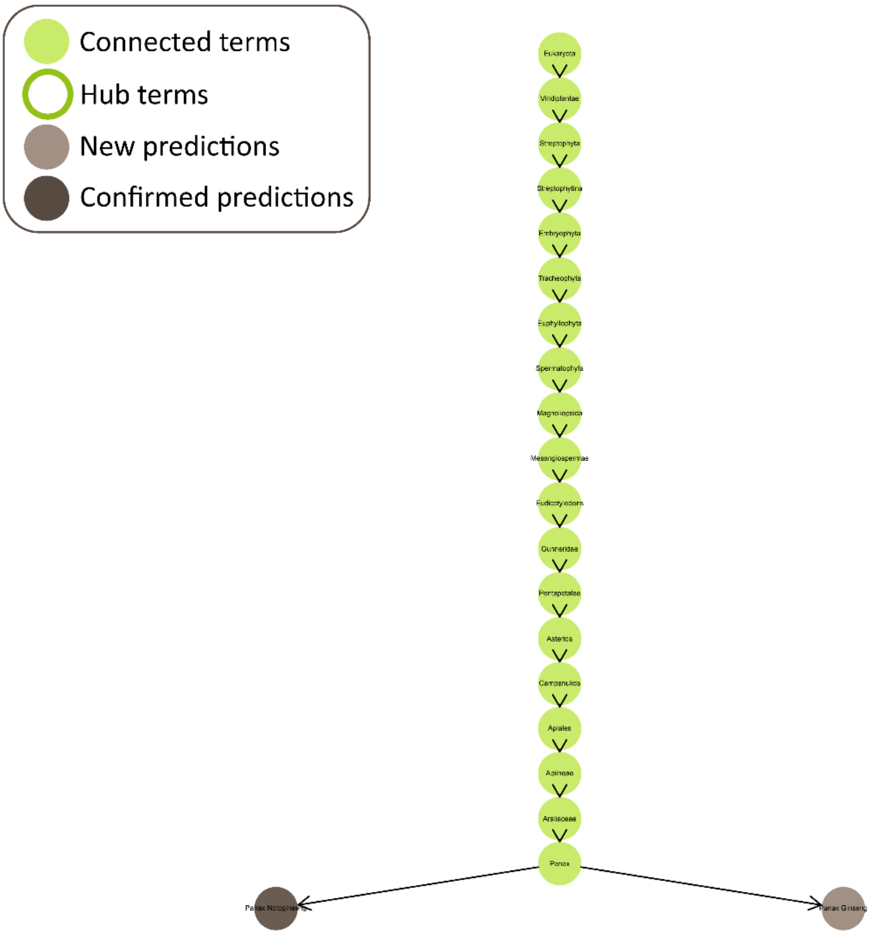

**Table S3**. Comparison of (un)validated CCT phenotypes associated with *Panax Notoginseng* constituents vs. the similar *Panax Ginseng*

| **Phenotype name** | **Also associated with 'Panax Ginseng'** | **Number of connected compounds (out of 46)** | **Connected compound name(s)** |
|---|---|---|---|
| Mitral valve prolapse | Yes | 4 | 2,6-di-tert-butyl-4-methylphenol, (-)-beta-elemene, delta-guaiene, alpha-guaiene |
| Histiocytoid cardiomyopathy | Yes | 2 | protopanaxadiol, panaxatriol |
| Overriding aorta | Yes | 2 | protopanaxadiol, panaxatriol |
| Shock | Yes | 1 | ginsenoside |
| Hypertensive retinopathy | Yes | 1 | protopanaxadiol |
| Vasospasm | Yes | 1 | protopanaxadiol |
| Vascular dilatation | No | 1 | 2,6-di-tert-butyl-4-methylphenol |
| Ventricular preexcitation | No | 0 | |
| Effort-induced polymorphic ventricular tachycardia | No | 0 | |
| Dilated cardiomyopathy | No | 0 | |
| Capillary fragility | No | 0 | |
| Cardiomegaly | No | 0 | |
| Supravalvular aortic stenosis | No | 0 | |
| Hypoplastic left heart | No | 0 | |
| Ventricular septal defect | No | 0 | |

*Note.* Phenotypes are validated by being both associated with *Panax ginseng* and sharing connected compounds.

**Figure S8.** NCBI ontology tree of herb similarity for validation of new CCT diseases using *Notoginseng*

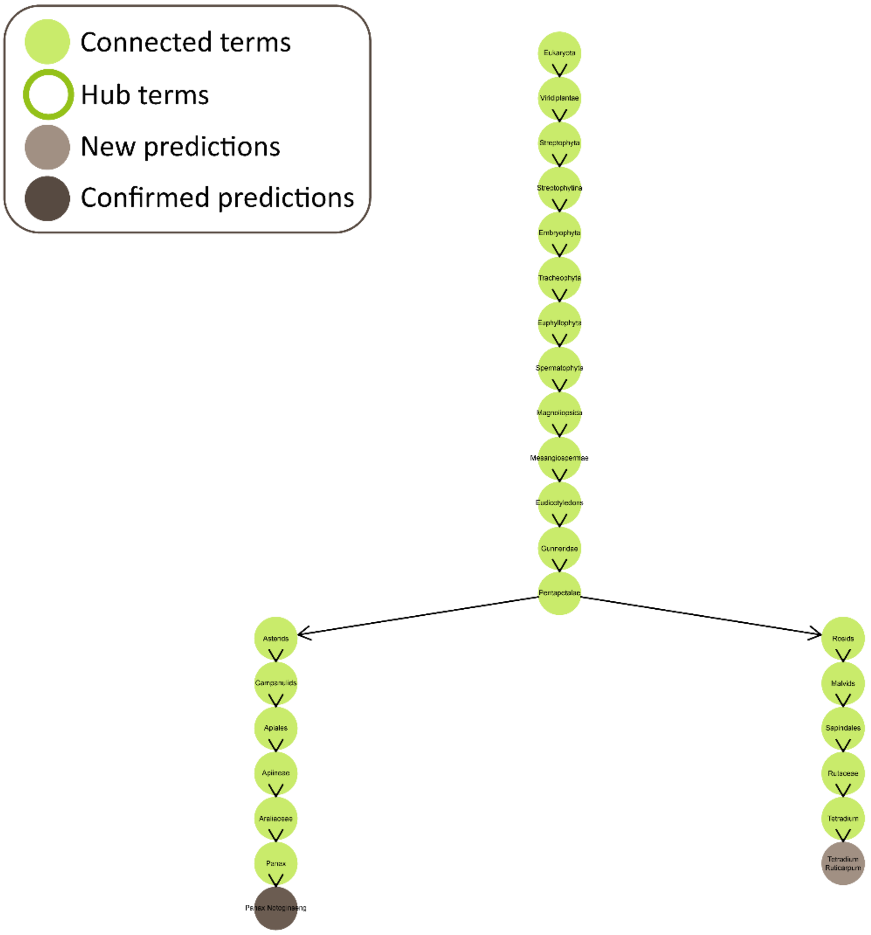
