## Supplementary PTS Quantitative Report for "Herb-paths, a network and statistical model to explore health-beneficial effects of herbs and herbal constituents"

**Quantitative report of components in panaxatriol saponins (PTS) by HPLC**

**1. Method**

1.1 Sample preparation

*Test solution*: Add 0.500 g of PTS (Lot No.: 200908, was provided by Chengdu Huasun Technology Group) into a 25 mL volumetric flask, dissolve with 20 mL of 15% acetonitrile aqueous solution, and sonicate for 30 min. After cooling, dilute to 25 mL with the same solvent, shake well, and filter through a 0.45 μm microporous membrane, and take the subsequent filtrate for measurement.

*Reference solution*: Precisely weigh 11 reference substances (see details in Table 1), respectively, dissolve with methanol to prepare the concentration whose peak area is close to the sample under the same injection volume, mix and shake well, filter through a 0.45 μm microporous membrane, and take the subsequent filtrate for measurement.

**Table 1 Information of reference substances**

| Name of reference substances | Batch no. | Supplier |
|---|---|---|
| Notoginsenoside $R_1$ | 110745-201820 | Chinese National Institutes for Food and Drug Control |
| Ginsenoside $Rg_1$ | 110703-201832 | |
| Ginsenoside Re | 110754-201123 | |
| Ginsenoside $Rb_1$ | 110704-201625 | |
| Notoginsenoside $R_2$ | PS010065 | Lovan Research chemicals |
| R-Notoginsenoside $R_2$ | PS210406-17 | |
| Ginsenoside $Rh_1$ | PS010067 | |
| R-Ginsenoside $Rh_1$ | PS010068 | |
| Ginsenoside $F_2$ | PS020390 | |
| Ginsenoside $Rk_3$ | PS010050 | |
| Ginsenoside $Rh_4$ | PCS-200923 | Chengdu herbsubstance Biotech |

1.2 Chromatographic Conditions

Chromatographic column: stainless steel column, with octadecylsilane bonded silica gel as filler (4.6 mm× 0.25 m, 5 μm); mobile phase: acetonitrile(A)- aqueous solution(B), the gradient elution program is shown in table 2; detection wavelength: 210 nm; column temperature: 30° C; flow rate : 1.0 mL/min; injection : 20 μL.

**Table 2 gradient elution program**

| Time (min) | A（% *V/V*） | B（% *V/V*） |
|---|---|---|
| 0～5 | 15 | 85 |
| 5~43 | 15→25 | 85→75 |
| 43~55 | 25→35 | 75→65 |
| 55~60 | 35→40 | 65→60 |
| 60~70 | 40→95 | 60→5 |
| 70~75 | 95→15 | 5→85 |
| 75~80 | 15 | 85 |

1.3 Samples detection

Take the PTS test solution, adopt the above-mentioned chromatographic conditions, and measure each batch with double samples and double needles. The content of each component was calculated by one-point external standard method, and the average value was taken.

The formula for calculating the content is:

$$\mathrm{Content\%} = \frac{\mathrm{A} \times \mathrm{p} \times \mathrm{C}\ (\mathrm{mg/mL})}{\mathrm{B} \times \mathrm{D}(\mathrm{mg/mL})} \times 100\%$$

A: Test solution peak area; B: Reference solution peak area; C: Concentration of reference substance; D: Sample concentration; p: percentage content of reference substance.

### 2 Results

The total content of 11 components in PTS is 88.75%, details are shown in Table 3 and Figure 1.

**Table 3 quantification date of 11 components in PTS**

| Peak NO. | retention time | Name | Formula | CAS NO. | content（%） |
|---|---|---|---|---|---|
| **1** | 34.585 | Notoginsenoside $R_1$ | $C_{47}H_{80}O_{18}$ | 80418-24-2 | 16.44 |
| **2** | 38.372 | Ginsenoside $Rg_1$ | $C_{42}H_{72}O_{14}$ | 22427-39-0 | 57.35 |
| **3** | 39.498 | Ginsenoside Re | $C_{48}H_{82}O_{18}$ | 52286-59-6 | 8.05 |
| **4** | 57.045 | Ginsenoside $Rb_1$ | $C_{54}H_{92}O_{23}$ | 41753-43-9 | 3.02 |
| **5** | 57.540 | Notoginsenoside $R_2$ | $C_{41}H_{70}O_{13}$ | 80418-25-3 | 0.63 |
| **6** | 58.604 | R-Notoginsenoside $R_2$ | $C_{41}H_{69}O_{13}$ | 948046-15-9 | 0.09 |
| **7** | 59.450 | Ginsenoside $Rh_1$ | $C_{36}H_{62}O_9$ | 63223-86-9 | 0.63 |
| **8** | 60.183 | R-Ginsenoside $Rh_1$ | $C_{36}H_{62}O_9$ | 80952-71-2 | 0.33 |
| **9** | 65.305 | Ginsenoside $F_2$ | $C_{42}H_{72}O_{13}$ | 62025-49-4 | 1.16 |
| **10** | 65.969 | Ginsenoside $Rk_3$ | $C_{36}H_{60}O_8$ | 364779-15-7 | 0.41 |
| **11** | 66.243 | Ginsenoside $Rh_4$ | $C_{36}H_{60}O_8$ | 174721-08-5 | 0.64 |

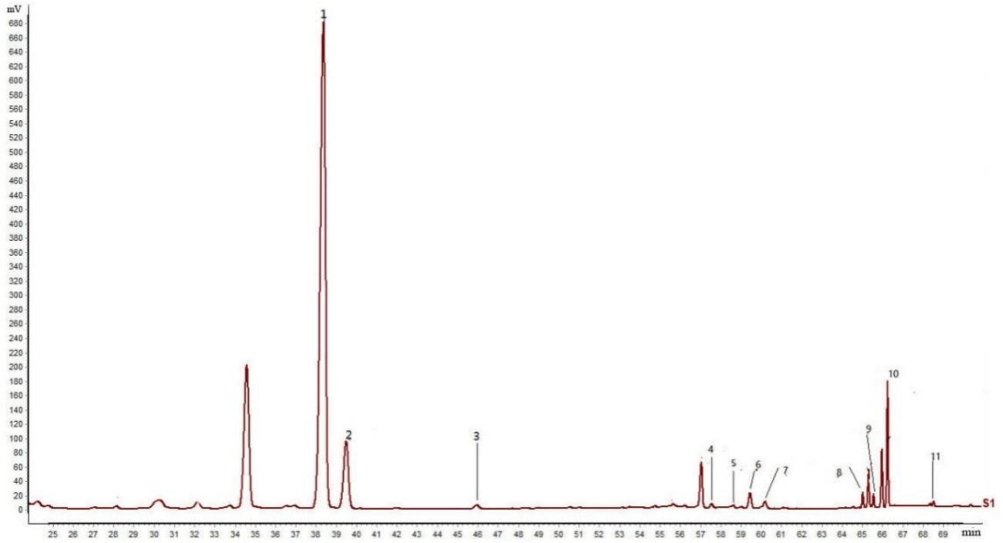


**Figure 1. HPLC chromatogram of PTS quantification**
