## Supplementary PTS Identification Report for "Herb-paths, a network and statistical model to explore health-beneficial effects of herbs and herbal constituents"

**Identification report of components in panaxatriol saponins (PTS) by UPLC-Q-TOF/MS**

**1. Method**

1.1 Sample preparation

Add 20 mg PTS (Lot No.: 210118, was provided by Chengdu Huasun Technology Group) into a centrifuge tube, dissolve with 1 mL methanol and weigh, ultrasonicate (power: 300 W, frequency: 40 kHz) for 30 min, let it cool to room temperature, weigh it again, complement the loss weight by same solvent, shake well, and centrifuge (12,000 rpm/min) for 5 min, and use the supernatant.

1.2 Test Conditions

1.2.1 Chromatographic Conditions

Chromatographic column: stainless steel column, with octadecylsilane bonded silica gel as filler (2.1 × 100 mm, 1.8 μm); mobile phase: 0.1% formic acid aqueous solution (A)-0.1% formic acid in acetonitrile (B), the gradient elution program is shown in table

1; detection wavelength: 190-400 nm; column temperature: 30° C; flow rate: 0.3 mL/min; injection: 1 μL.

**Table 1 gradient elution program**

| Time (min) | A（% *V/V*） | B（% *V/V*） |
|---|---|---|
| 0~3 | 92 | 8 |
| 3~8 | 95→85 | 8→15 |
| 8~22 | 85→80 | 15→20 |
| 22~30 | 80 | 20 |
| 30~37 | 80→72 | 20→28 |
| 37~52 | 72→55 | 28→45 |
| 52~55 | 55→5 | 45→95 |
| 55~57 | 5 | 95 |
| 57~57.1 | 5→92 | 95→8 |
| 57.1~61 | 92 | 8 |

1.2.2 Mass spectrometry conditions

ESI IDA mode was used to collect data; detection mode: Negative/Positive ion mode; the mass parameters are shown in Table 2.

**Table 2 Mass parameters of Sciex Triple TOF**

| MS parameters | parameter value | MS/MS parameters | parameter value |
|---|---|---|---|
| TOF mass range | 50～1700 | MS/MS mass range | 50～1250 |
| Ion Source Gas 1(psi) | 50 | Declustering Potential(V) | 100 |
| Ion Source Gas 2(psi) | 50 | Collision Energy(eV) | ±40 |
| Curtain Gas(psi) | 35 | Collision Energy Spread(eV) | 20 |
| Ion Spray Voltage Floating (V) | -4500/5000 | Ion Release Delay(ms) | 30 |
| Ion Source Temperature (°C) | 500 | Ion Release Width(ms) | 15 |
| Declustering Potential(V) | 100 | — | — |
| Collision Energy(eV) | 10 | — | — |

### 2. Analyses

Data acquisition software: Analyst TF 1.7.1; data analysis software: Peak View 1.2; database: Natural Products HR-MS/MS Spectra Library 1.0.

### 3. Results

The sample of PTS was analyzed by method of ultra-performance liquid chromatography-high-resolution mass spectrometry (UPLC-Q-TOF/MS).

As shown in Figures 1-2, there are 85 peaks in the UPLC-MS spectrum, 81 candidate components were identified (see details in Table 3), and the peaks numbered 23, 49, 53 and 85 were not identified, according to the multi-level mass spectrometry information of the sample, combined with the natural product high-resolution mass spectrometry database and related literatures.

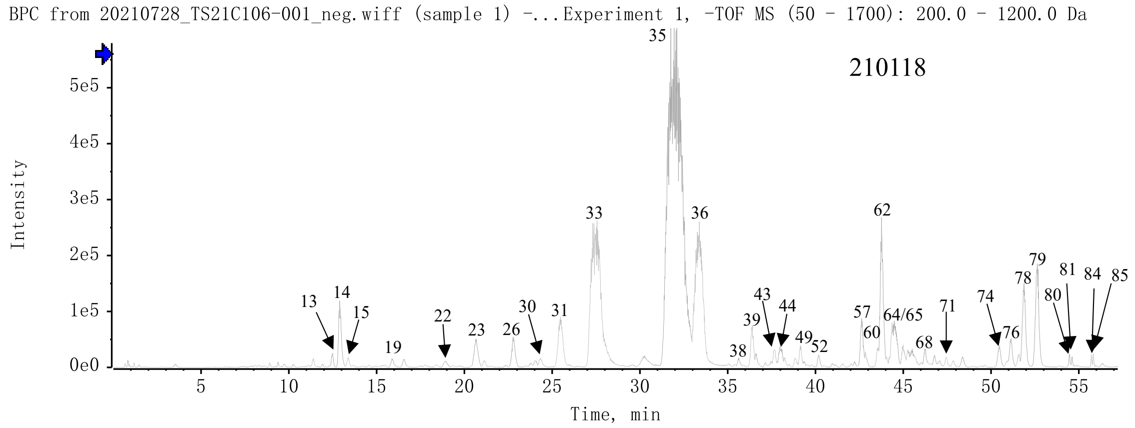


**Figure 1. UPLC-HRMS Base Peak Ion Chromatogram (BPC) - Negative ion mode**

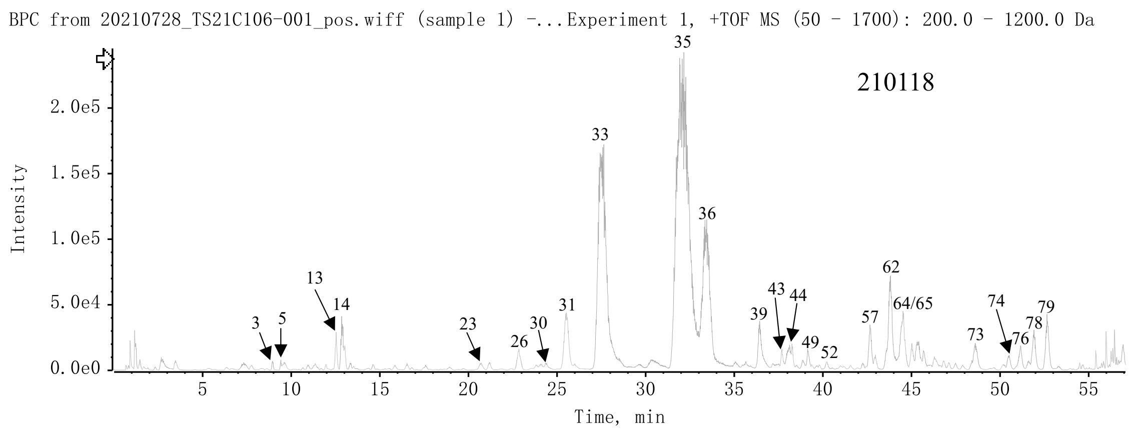


**Figure 2. UPLC-HRMS BPC - Positive ion mode**

**Table 3 Identification results of target components of PTS**

| Peak no. | Retention time | Adduct ion | Actual value （m/z） | Theoretical value （m/z） | ppm | Formula | molecular weight | CAS | Name | MS/MS | Database | Reference |
|---|---|---|---|---|---|---|---|---|---|---|---|---|
| **1** | 3.51 | $[M+H]^+$ | 205.0968 | 205.0972 | -1.7 | $C_{11}H_{12}N_2O_2$ | 204.23 | 73-22-3 | L-tryptophan | 188.0718;146.0602;118.0648 | √ | |
| **2** | 8.33 | $[M+FA-H]^-$ | 447.1505 | 447.1508 | -0.7 | $C_{18}H_{26}O_{10}$ | 402.39 | 2131787-98-7 | 3-Methylphenyl 6-O-β-D-xylopyranosyl-β-D-glucopyranoside | 401.1468;269.1015;161.0482 | | [1] |
| **3** | 8.93 | $[M+FA-H]^-$ | 879.4933 | 879.4933 | -2.9 | $C_{42}H_{74}O_{16}$ | 835.03 | 193977-11-6 | Notoginsenoside J | 833.4947;671.4483;653.4324;179.0557 | | [2] |
| **4** | 9.37 | $[M+FA-H]^-$ | 1011.5379 | 1011.5381 | -0.2 | $C_{47}H_{82}O_{20}$ | 967.14 | 1133882-74-2 | (3β,6β,12β)-20-(β-D-Glucopyranosyloxy)-3,12,24,25-tetrahydroxydammaran-6-yl 6-O-β-D-xylopyranosyl-β-D-glucopyranoside | 1011.5449;965.5318;785.4673;653.4279 | | [3] |
| **5** | 9.39 | $[M+FA-H]^-$ | 879.4981 | 879.4959 | 2.5 | $C_{42}H_{74}O_{16}$ | 835.03 | 226713-28-6 | Vinaginsenoside R22 | 833.4875;671.4316;653.4235;179.0560 | | [4] |
| **6** | 9.77 | [M+FA | 1011.53 | 1011.5381 | 1.5 | $C_{47}H_{82}O_{20}$ | 967.14 | 1394905-99-7 | (3β,6α,12β)-3,12,24,25-Tetrahydroxy-20-(β- | 1011.5394;9 | | [5] |

| | | | | | | | | | | | | |
|---|---|---|---|---|---|---|---|---|---|---|---|---|
| | | -H]- | 97 | | | | | | D-glucopyranosyloxy) dammaran-6-yl O-pentosyl-β-D-glucopyranoside | 65.5343;785.4643;653.4287 | | |
| **7** | 10.26 | [M+FA-H]- | 503.1769 | 503.1770 | -0.2 | $C_{22}H_{32}O_{13}$ | 504.48 | 1347015-00-2 | Isoconiferoside | 503.1781;341.1259;221.0661;179.0703;161.0458 | | [6] |
| **8** | 10.30 | [M+FA-H]- | 1025.5591 | 1025.5538 | 5.2 | $C_{48}H_{84}O_{20}$ | 981.17 | 1133882-75-3 | (3β, 6β, 12β) - 20- (β- D- Glucopyranosyloxy) - 3, 12, 24, 25- tetrahydroxydammaran - 6- yl 6- O- (6- deoxy- α- L- mannopyranosyl) - β- D- glucopyranoside | 1025.5561;979.5453;799.4806;653.4327 | | [3] |
| **9** | 11.40 | [M-H]- | 625.1406 | 625.1410 | -0.7 | $C_{27}H_{30}O_{17}$ | 626.52 | 18609-17-1 | Baimaside | 625.1377;300.0250;271.0226;255.0281 | √ | |
| **10** | 11.66 | [M+FA-H]- | 847.4753 | 847.4697 | 6.6 | $C_{41}H_{70}O_{15}$ | 802.99 | 942586-98-3 | Floralginsenoside C | 847.4727;801.4704;639.4040;477.3594 | √ | |
| **11** | 11.92 | [M+FA-H]- | 993.5275 | 993.5276 | -0.1 | $C_{47}H_{80}O_{19}$ | 949.13 | 193976-69-1 | Notoginsenoside H | 947.5259;767.4608;635.4136 | | [2] |
| **12** | 11.94 | [M+FA-H]- | 861.4888 | 861.4853 | 4.0 | $C_{42}H_{72}O_{15}$ | 817.01 | 1801723-21-6 | Notoginsenoside SP1 | 861.4841;815.4794;653. | | [7] |

| | | | | | | | | | | | | |
|---|---|---|---|---|---|---|---|---|---|---|---|---|
| | | | | | | | | | | 4325;635.41 41;179.0556 | | |
| **13** | 12.47 | [M+FA -H]- | 861.483 3 | 861.4853 | -2.4 | $C_{42}H_{72}O_{15}$ | 817.01 | 1801723-22-7 | Notoginsenoside SP2 | 815.4803;65 3.4241;635. 4152;553.31 97;491.3753 | | [7] |
| **14** | 12.92 | [M+FA -H]- | 595.133 3 | 595.1305 | 4.8 | $C_{26}H_{28}O_{16}$ | 596.49 | 83048-35-5 | Quercetin 3-O-sambubioside | 595.1323;30 0.0263;271. 0227;255.02 74 | | [8] |
| **15** | 13.37 | [M-H]- | 609.145 7 | 609.1461 | -0.7 | $C_{27}H_{30}O_{16}$ | 610.52 | 25615-14-9 | Kaempferol 3,7-diglucoside | 609.1454;28 5.0398;225. 0352 | | [2] |
| **16** | 14.56 | [M+FA -H]- | 877.477 6 | 877.4802 | -3.0 | $C_{42}H_{72}O_{16}$ | 833.01 | 942586-97-2 | Floralginsenoside B | 877.4859;83 1.4758;651. 4123;179.05 63;161.0466 | | [9] |
| **17** | 15.37 | [M-H]- | 463.087 8 | 463.0882 | -0.9 | $C_{21}H_{20}O_{12}$ | 464.38 | 482-36-0 | Hyperoside | 463.0852;30 0.0252;271. 0219;255.02 85 | √ | |
| **18** | 15.50 | [M+FA -H]- | 859.473 9 | 859.4697 | 4.9 | $C_{42}H_{70}O_{15}$ | 815.00 | 340270-89-5 | Vina- ginsenoside R25 | 813.4579;63 3.4048;471. 3407;179.05 58 | | [4] |

| | | | | | | | | | | | | |
|---|---|---|---|---|---|---|---|---|---|---|---|---|
| **19** | 15.92 | $[M-H]^-$ | 579.1377 | 579.1355 | 3.7 | $C_{26}H_{28}O_{15}$ | 580.49 | 27661-51-4 | Kaempferol 3-O-sambubioside | 579.1390;285.0325;255.0302;227.0356 | | [8] |
| **20** | 15.94 | $[M-H]^-$ | 463.0862 | 463.0882 | -4.3 | $C_{21}H_{20}O_{12}$ | 464.38 | 482-35-9 | Isoquercetin | 463.0885;300.0262;271.0213;243.0284 | √ | |
| **21** | 17.55 | $[M+FA-H]^-$ | 859.4692 | 859.4697 | -0.6 | $C_{42}H_{70}O_{15}$ | 815.00 | 1135445-37-2 | (3β, 6β, 12β) - 6, 20- Bis(β- D-glucopyranosyloxy) - 3, 12-dihydroxydammar- 25- en- 24- one | 859.4719;813.4712;651.4105;633.3933;179.0573 | | [10] |
| **22** | 18.93 | $[M+FA-H]^-$ | 517.1950 | 517.1927 | 4.5 | $C_{22}H_{32}O_{11}$ | 472.48 | 138772-01-7 | Eugenol rutinoside | 471.1880;325.1283;163.0760 | | [11] |
| **23** | 20.66 | $[M+FA-H]^-$ | 469.2297 | 469.2291 | 1.4 | $C_{19}H_{36}O_{10}$ | 424.48 | / | / | 423.2254;277.1642;161.0442 | | |
| **24** | 21.09 | $[M+FA-H]^-$ | 1139.5905 | 1139.5855 | 4.4 | $C_{53}H_{90}O_{23}$ | 1095.27 | 109150-50-7 | Gypenoside LXIX | 1139.5853;1093.5678;961.5181 | | [12] |
| **25** | 22.36 | $[M+FA-H]^-$ | 1139.5924 | 1139.5855 | 6.1 | $C_{53}H_{90}O_{23}$ | 1095.27 | 105214-48-0 | Gypenoside LVI | 1139.5970;1093.5791;961.5233 | | [13] |
| **26** | 22.78 | $[M+FA-H]^-$ | 1007.5493 | 1007.5432 | 6.0 | $C_{48}H_{82}O_{19}$ | 963.15 | 350586-56-0 | Notoginsenoside N | 1007.5447;961.5375;799 | | [12] |

| | | | | | | | | | | | | |
|---|---|---|---|---|---|---|---|---|---|---|---|---|
| | | | | | | | | | | .4866;637.4343 | | |
| **27** | 23.48 | [M+FA-H]$^-$ | 1139.5876 | 1139.5855 | 1.8 | $C_{53}H_{90}O_{23}$ | 1095.27 | 1179351-11-1 | Floranotoginsenoside A | 1139.6428;1093.5776;931.5191 | | [9] |
| **28** | 23.78 | [M+FA-H]$^-$ | 1007.5413 | 1007.5432 | -1.9 | $C_{48}H_{82}O_{19}$ | 963.15 | 156042-22-7 | Vinaginsenoside R8 | 1007.5421;961.5359;799.4834 | √ | |
| **29** | 24.10 | [M+FA-2H]$^{2-}$ | 599.3004 | 599.2997 | 1.2 | $C_{54}H_{92}O_{23}$ | 1109.29 | 94705-68-7 | Gypenoside XLIII | 1107.6018;945.5346;783.4721;637.4272;553.2986 | | [13] |
| **30** | 24.33 | [M+FA-H]$^-$ | 1007.5473 | 1007.5432 | 4.0 | $C_{48}H_{82}O_{19}$ | 963.15 | 394246-74-3 | Notoginsenoside M | 1007.5468;961.5402;799.5129;637.4333 | | [12] |
| **31** | 25.51 | [M+FA-H]$^-$ | 1007.5449 | 1007.5432 | 1.7 | $C_{48}H_{82}O_{19}$ | 963.15 | 87741-78-4 | Notoginsenoside-R6 | 961.5440;799.5162;637.4274;475.3776 | | [6] |
| **32** | 25.99 | [M+FA-H]$^-$ | 977.5328 | 977.5327 | 0.1 | $C_{47}H_{80}O_{18}$ | 933.13 | 1255210-79-7 | Ginsenoside Re4 | 977.5330;931.5338;799.4775;637.4353 | | [14] |
| **33** | 27.49 | [M+FA | 977.535 | 977.5327 | 2.9 | $C_{47}H_{80}O_{18}$ | 933.13 | 80418-24-2 | Notoginsenoside R1 | 931.5275;79 | √ | |

| | | | | | | | | | | | | |
|---|---|---|---|---|---|---|---|---|---|---|---|---|
| | | -H]$^-$ | 5 | | | | | | | 9.4858;637.4329;475.3779 | | |
| **34** | 28.45 | [M+FA-H]$^-$ | 1007.5474 | 1007.5432 | 4.1 | $C_{48}H_{82}O_{19}$ | 963.15 | 68406-27-9 | 20-Glucoginsenoside Rf | 1007.5506;961.5411;799.4879;637.4437 | | [15] |
| **35** | 31.86 | [M+FA-H]$^-$ | 845.4912 | 845.4904 | 0.9 | $C_{42}H_{72}O_{14}$ | 801.01 | 22427-39-0 | Ginsenoside Rg1 | 845.4892;799.4827;637.4303;475.3782 | √ | |
| **36** | 33.43 | [M+FA-H]$^-$ | 991.5505 | 991.5483 | 2.2 | $C_{48}H_{82}O_{18}$ | 947.15 | 52286-59-6 | Ginsenoside Re | 945.5388;783.4882;637.4290;475.3791 | √ | |
| **37** | 35.42 | [M+FA-H]$^-$ | 1025.5561 | 1025.5538 | 2.2 | $C_{48}H_{84}O_{20}$ | 981.17 | 1133882-74-2 | (3β,6β,12β)-20-(β- D- Glucopyranosyloxy)-3,12,25-trihydroxydammaran-6-yl 2-O-β-D-glucopyranosyl-β-D-glucopyranoside | 1025.5483;979.5440;817.5391;799.4811;637.4405 | | [3] |
| **38** | 35.66 | [M+FA-2H]$^{2-}$ | 616.3044 | 616.3024 | 3.2 | $C_{54}H_{94}O_{25}$ | 1143.31 | 454686-08-9 | Quinquenoside L16 | 1141.6022;979.5428;817.4983;799.4845;570.2957 | | [5] |

| | | | | | | | | | | | | |
|---|---|---|---|---|---|---|---|---|---|---|---|---|
| **39** | 36.40 | [M+FA-H]$^-$ | 887.5010 | 887.5010 | 0.0 | $C_{44}H_{74}O_{15}$ | 843.05 | 163403-91-6 | Yesanchinoside D | 841.5001;799.4786;781.4786;637.4330;475.3844 | √ | |
| **40** | 36.63 | [M+FA-H]$^-$ | 1169.6034 | 1169.5961 | 6.3 | $C_{54}H_{92}O_{24}$ | 1125.29 | 193895-21-5 | Notoginsenoside A | 1123.5965;961.5549;799.4999 | | [12] |
| **41** | 37.15 | [M+FA-H]$^-$ | 1025.5591 | 1025.5538 | 5.2 | $C_{48}H_{84}O_{20}$ | 981.17 | 156398-72-0 | Vinaginsenoside R13 | 979.5517;817.4929;799.4863;637.4201 | | [4] |
| **42** | 37.34 | [M+FA-H]$^-$ | 815.4835 | 815.4798 | 4.5 | $C_{41}H_{70}O_{13}$ | 770.99 | 98474-75-0 | Pseudoginsenoside RT3 | 815.4802;769.4712;607.4221;475.3821 | | [15] |
| **43** | 37.69 | [M+FA-H]$^-$ | 1005.5286 | 1005.5276 | 1.0 | $C_{48}H_{80}O_{19}$ | 961.14 | 223710-06-3 | Ginsenoside III | 959.5236;797.4678;635.4097;473.3617 | | [16] |
| **44** | 38.04 | [M+FA-H]$^-$ | 887.5018 | 887.5010 | 0.9 | $C_{44}H_{74}O_{15}$ | 843.05 | / | Yesanchinoside D isomer | 887.5054;841.4984;799.4849;781.4746;637.4356 | | [15] |
| **45** | 38.20 | [M+FA-H]$^-$ | 815.4799 | 815.4798 | 0.1 | $C_{41}H_{70}O_{13}$ | 770.99 | 189513-26-6 | Ginsenoside F5 | 815.4846;769.4781;637.4344;619.43 | √ | |

| | | | | | | | | | | | | |
|---|---|---|---|---|---|---|---|---|---|---|---|---|
| | | | | | | | | | | 10 | | |
| **46** | 38.25 | $[M+FA-2H]^{2-}$ | 607.2986 | 607.2971 | 2.4 | $C_{54}H_{92}O_{24}$ | 1125.29 | 2172820-99-2 | Notoginsenoside Fh7 | 1123.5955;961.5587;781.4777;561.2954;179.0563 | | [17] |
| **47** | 38.52 | $[M+FA-2H]^{2-}$ | 658.3161 | 658.3130 | 4.7 | $C_{58}H_{98}O_{27}$ | 1227.38 | 2172820-95-8 | Notoginsenoside Fh3 | 1225.6223;1093.5413;961.5208;799.4891;612.3088 | | [17] |
| **48** | 38.84 | $[M+FA-H]^{-}$ | 815.4807 | 815.4798 | 1.0 | $C_{41}H_{70}O_{13}$ | 770.99 | 62025-50-7 | Ginsenoside F3 | 815.4836;769.4804;637.4349;475.3819 | √ | |
| **49** | 39.16 | $[M+FA-H]^{-}$ | 887.5026 | 887.5010 | 1.8 | $C_{44}H_{74}O_{15}$ | 843.05 | / | / | 887.4992;841.4929;799.4829;781.4727;637.4270 | | |
| **50** | 39.35 | $[M+FA-2H]^{2-}$ | 606.2895 | 606.2893 | 0.3 | $C_{54}H_{90}O_{24}$ | 1123.28 | 193895-26-0 | Notoginsenoside B | 1121.5798;959.5231;797.4707;635.4157;560.2891 | | [15] |

| | | | | | | | | | | | | |
|---|---|---|---|---|---|---|---|---|---|---|---|---|
| **51** | 39.80 | [M+FA -H]- | 843.4736 | 843.4748 | -1.4 | $C_{42}H_{70}O_{14}$ | 799.00 | 1614215-08-5 | (6β, 12β) - 6, 20- Bis(β- D-glucopyranosyloxy) - 12- hydroxydammar-24- en- 3- one | 843.4800;797.4714;635.4134;617.4073 | | [10] |
| **52** | 40.19 | [M+FA -H]- | 1007.5454 | 1007.5432 | 1.3 | $C_{48}H_{82}O_{19}$ | 963.15 | 156009-80-2 | Vina-ginsenoside R4 | 1007.5453;961.5337;799.4794;637.4294 | √ | |
| **53** | 40.97 | [M+FA -H]- | 887.5019 | 887.5010 | 1.0 | $C_{44}H_{74}O_{15}$ | 843.05 | / | / | 887.5051;841.5007;799.4874;781.4779;637.4344 | / | / |
| **54** | 41.57 | [M+FA -H]- | 845.4908 | 845.4904 | 0.5 | $C_{42}H_{72}O_{14}$ | 801.01 | 52286-58-5 | Ginsenoside Rf | 845.4903;799.4833;637.4314;475.3836 | √ | |
| **55** | 42.05 | $[M+FA-2H]^{2-}$ | 731.3424 | 731.3419 | 0.6 | $C_{64}H_{108}O_{31}$ | 1373.52 | 193895-50-0 | Notoginsenoside D | 1077.5459;945.5257;783.4904;685.3323 | | [18] |
| **56** | 42.23 | [M+FA -H]- | 845.4932 | 845.4904 | 3.3 | $C_{42}H_{72}O_{14}$ | 801.01 | 94987-09-4 | Gypenoside L | 845.5031;799.4919;637.4405;475.3789 | | [13] |
| **57** | 42.63 | [M+FA -H]- | 815.4775 | 815.4798 | -2.9 | $C_{41}H_{70}O_{13}$ | 770.99 | 80418-25-3 | 20(S)-Notoginsenoside R2 | 769.4711;637.4283;475. | √ | |

| | | | | | | | | | | | | |
|---|---|---|---|---|---|---|---|---|---|---|---|---|
| | | | | | | | | | | 3768 | | |
| **58** | 42.84 | [M+FA-2H]$^{2-}$ | 665.3213 | 665.3208 | 0.7 | $C_{59}H_{100}O_{27}$ | 1241.41 | 137348-15-3 | Chikusetsusaponin VI | 1239.6371;945.5376;783.4875;619.3170 | | [12] |
| **59** | 43.54 | [M+FA-2H]$^{2-}$ | 665.3227 | 665.3206 | 2.9 | $C_{59}H_{100}O_{27}$ | 1241.41 | 88100-04-3 | Notoginsenoside Fa | 1239.6164;1107.5616;945.5564;783.4729;619.3157 | √ | |
| **60** | 43.55 | [M+FA-H]$^{-}$ | 815.4760 | 815.4798 | -4.7 | $C_{41}H_{70}O_{13}$ | 770.99 | 948046-15-9 | 20(R)-Notoginsenoside R2 | 769.4706;637.4309;475.3782 | √ | |
| **61** | 43.74 | [M+Cl]$^{-}$ | 819.4682 | 819.4667 | 1.8 | $C_{42}H_{72}O_{13}$ | 785.01 | 80952-72-3 | 20(S)-Ginsenoside Rg2 | 819.4647;783.4879;637.4249;475.3761 | √ | |
| **62** | 43.75 | [M+FA-H]$^{-}$ | 683.4395 | 683.4376 | 2.8 | $C_{36}H_{62}O_{9}$ | 638.87 | 63223-86-9 | 20(S)-Ginsenoside Rh1 | 683.4399;637.4364;475.3803 | √ | |
| **63** | 44.05 | [M+FA-H]$^{-}$ | 829.4983 | 829.4955 | 3.4 | $C_{42}H_{72}O_{13}$ | 785.01 | 80952-72-3 | 20(R)-Ginsenoside Rg2 | 829.4937;783.4917;637.4352;475.3781 | | [6] |
| **64** | 44.36 | [M+FA-H]$^{-}$ | 683.4387 | 683.4376 | 1.6 | $C_{36}H_{62}O_{9}$ | 638.87 | 80952-71-2 | 20(R)-Ginsenoside Rh1 | 683.4377;637.4326;475. | √ | |

| | | | | | | | | | | | | |
|---|---|---|---|---|---|---|---|---|---|---|---|---|
| | | | | | | | | | | 3779 | | |
| **65** | 44.52 | [M+FA -H]- | 1153.6007 | 1153.6011 | -0.4 | $C_{54}H_{92}O_{23}$ | 1109.29 | 41753-43-9 | Ginsenoside Rb1 | 1107.5944;945.5394;783.4920;621.4467 | √ | |
| **66** | 45.00 | [M-H]- | 1193.5971 | 1193.5961 | 0.9 | $C_{57}H_{94}O_{26}$ | 1195.34 | 88140-34-5 | Malonyl-ginsenoside Rb1 | 1149.6107;1107.5973;945.5446;783.5018;621.4322 | | [6] |
| **67** | 45.00 | [M+FA -2H]2- | 620.3049 | 620.3050 | -0.1 | $C_{56}H_{94}O_{24}$ | 1151.33 | 88140-34-5 | Quinquenoside R1 | 1149.6020;1107.6102;945.5449;783.4820;574.3012 | | [4] |
| **68** | 46.26 | [M+FA -H]- | 683.4386 | 683.4376 | 1.5 | $C_{36}H_{62}O_9$ | 638.87 | 53963-43-2 | Ginsenoside F1 | 683.4362;637.4299;475.3788 | √ | |
| **69** | 46.80 | [M-H]- | 925.4788 | 925.4802 | -1.6 | $C_{47}H_{74}O_{18}$ | 927.08 | 7518-22-1 | Chikusetsusaponin IV | 925.4802;775.4376;613.3764;569.3839 | | [6] |
| **70** | 47.13 | [M+FA -H]- | 857.4906 | 857.4904 | 0.2 | $C_{43}H_{72}O_{14}$ | 813.02 | 1702363-34-5 | 20S- Sanchirhinoside A2 | 811.4840;769.4556;679.4331;637.4394;475.3831 | | [5] |

| | | | | | | | | | | | | |
|---|---|---|---|---|---|---|---|---|---|---|---|---|
| **71** | 47.45 | [M+FA-H]- | 991.5474 | 991.5483 | -0.9 | $C_{48}H_{82}O_{18}$ | 947.15 | 52705-93-8 | Ginsenoside Rd | 991.5461;945.5394;783.4921;621.4374 | √ | |
| **72** | 47.86 | [M+FA-H]- | 725.4463 | 725.4325 | -4.4 | $C_{38}H_{64}O_{10}$ | 680.91 | 784213-07-6 | 6- acetate(3β, 6α, 12β) - 3, 12, 20-Trihydroxydammar- 24- en- 6- yl β- D-glucopyranoside | 725.4480;679.4429;637.4307;619.4307;475.3783 | | [19] |
| **73** | 48.39 | [M+FA-H]- | 725.4460 | 725.4482 | -3.0 | $C_{38}H_{64}O_{10}$ | 680.91 | 1133882-72-0 | (3β,6β,12β)-6-(Acetyloxy)-3,12-dihydroxydammar-24-en--20-yl β-D-glucopyranoside | 725.4482;679.4390;637.4241;619.4223;475.3806 | | [10] |
| **74** | 50.50 | [M+FA-H]- | 797.4691 | 797.4693 | -0.2 | $C_{41}H_{68}O_{12}$ | 752.97 | 1375079-47-2 | (3β,12β)-12-Hydroxy-6-(D-xylopyranosyloxy) dammara-20, 24(or 20(22) , 24)-dien-3-yl β-D-glucopyranoside | 797.4772;751.4658;619.4203 | | [10] |
| **75** | 50.94 | [M+FA-H]- | 811.4835 | 811.4849 | -1.8 | $C_{42}H_{70}O_{12}$ | 767.00 | 126223-28-7 | Ginsenoside Rg4 | 811.4883;765.4780;619.4215 | | [20] |
| **76** | 51.15 | [M+FA-H]- | 797.4677 | 797.4693 | -2.0 | $C_{41}H_{68}O_{12}$ | 752.97 | 1375079-55-2 | (3β,12β)-12-Hydroxy-3-(D-xylopyranosyloxy) dammara-20, 24(or 20(22) , 24) -dien-6-yl β-D-glucopyranoside | 797.4688;751.4617;619.4197 | | [10] |
| **77** | 51.58 | [M+FA-H]- | 811.4860 | 811.4849 | 1.3 | $C_{42}H_{70}O_{12}$ | 767.00 | 147419-93-0 | Ginsenoside Rg6 | 811.4878;765.4800;619.4230;601.4127;161.0454 | √ | |

| | | | | | | | | | | | | |
|---|---|---|---|---|---|---|---|---|---|---|---|---|
| **78** | 51.88 | [M+FA-H]- | 665.4278 | 665.4270 | 1.2 | $C_{36}H_{60}O_8$ | 620.86 | 364779-15-7 | Ginsenoside Rk3 | 665.4220;619.4177;161.0455 | √ | |
| **79** | 52.63 | [M+FA-H]- | 665.4285 | 665.4270 | 2.2 | $C_{36}H_{60}O_8$ | 620.86 | 174721-08-5 | Ginsenoside Rh4 | 665.4281;619.4243;161.0448 | √ | |
| **80** | 54.47 | [M+FA-H]- | 829.4951 | 829.4955 | -0.5 | $C_{42}H_{72}O_{13}$ | 785.01 | 62025-49-4 | Ginsenoside F2 | 829.4882;783.4850;621.4371;459.3857 | √ | |
| **81** | 54.62 | [M+FA-H]- | 829.4954 | 829.4955 | -0.1 | $C_{42}H_{72}O_{13}$ | 785.01 | 14197-60-5 | Ginsenoside Rg3 | 829.4995;783.4931;621.4359;459.3857 | √ | |
| **82** | 54.64 | [M+FA-H]- | 871.5061 | 871.5061 | 0.0 | $C_{44}H_{74}O_{14}$ | 827.05 | 203849-13-2 | 20(R)-Ginsenoside Rs3 | 825.5022;783.4900;765.4802;621.4353;459.3804 | | [21] |
| **83** | 54.76 | [M+FA-H]- | 871.5073 | 871.5061 | 1.4 | $C_{44}H_{74}O_{14}$ | 827.05 | 194861-70-6 | Ginsenoside Rs3 | 871.5084;825.5041;783.4889;621.4312;459.3860 | | [15] |
| **84** | 55.72 | [M+FA-H]- | 811.4868 | 811.4849 | 2.3 | $C_{42}H_{70}O_{12}$ | 767.00 | 41753-43-9 | Ginsenoside Rg5 | 811.4864;765.4790;603.4270;161.0457 | √ | |

| | | | | | | | | | | | | |
|---|---|---|---|---|---|---|---|---|---|---|---|---|
| **85** | 55.83 | [M+FA-H]$^-$ | 811.4857 | 811.4849 | 0.9 | $C_{42}H_{70}O_{12}$ | 767.00 | / | / | 811.4922;765.4797;603.4333;161.0480 | / | / |

**REFERENCES**


(1) Gu, C.-Z.; Qiao, Y.-J.; Wang, D.; Zhu, H.-T.; Yang, C.-R.; Xu, M.; Zhang, Y.-J. New triterpenoid saponins from the steaming treated roots of Panax notoginseng. *Nat. Prod. Res*. **2018**, *32*(3), 294-301. https://doi.org/10.1080/14786419.2017.1356833.

(2) Wang, L.; Wang, Y.; Tong, G.; Li, Y.; Lei, M.; Wu, H.; Wang, B.; Hu, R. Development of a novel UHPLC-UV combined with UHPLC-QTOF/MS fingerprint method for the comprehensive evaluation of nao-luo-xin-tong: multi-wavelength setting based on traditional chinese medicinal prescription composition. *Anal. Methods-UK*. **2019**, *11*(48), 6092-6102. https://doi.org/10.1039/C9AY01975H.

(3) Liu, Y.; Li, J.; He, J.; Abliz, Z.; Qu, J.; Yu, S.; Ma, S.; Liu, J.; Du, D. Identification of new trace triterpenoid saponins from the roots of Panax notoginseng by high-performance liquid chromatography coupled with electrospray ionization tandem mass spectrometry. *Rapid. Commun. Mass. SP*. **2009**, *23*(5), 667-679. https://doi.org/10.1002/rcm.3917.

(4) Do, T. T. T.; Nguyen, H. T. T.; Duong, Q. H. T.; Le, S. H.; Nguyen, P. T. V. Virtual screening of saponin derivatives targeting enzymes endothelial nitric oxide synthase and cytochrome P450 2E1. *Int. J. Pharm. Sci. Res*. 2019, *10*(1), 70-82. https://ijpsr.com/?action=download_pdf&postid=47734.

(5) Zhang, Y.; Han, L.-F.; Sakah, K.J.; Wu, Z.-Z.; Liu, L.-L.; Agyemang, K.; Gao, X.-M.; Wang, T. Bioactive protopanaxatriol type saponins isolated from the roots of Panax notoginseng (Burk.) F. H. Chen. *Molecules*. **2013**, *18*(9), 10352-10366. https://doi.org/10.3390/molecules180910352.

(6) Lin, H.; Zhu, H.; Tan, J.; Wang, C.; Dong, Q.; Wu, F.; Wang, H.; Liu, J.; Li, P.; Liu, J. Comprehensive Investigation on Metabolites of Wild-Simulated American Ginseng Root Based on Ultra-High Performance Liquid Chromatography-Quadrupole Time-of-Flight Mass Spectrometry. *J. Agric. Food. Chem*. **2019**, *67*(20), 5801-5819. https://doi.org/10.1021/acs.jafc.9b01581.

(7) Gu, C.-Z.; Lv, J.-J.; Zhang, X.-X.; Qiao, Y.-J.; Yan, H.; Li, Y.; Wang, D.; Zhu, H.-T.; Luo, H.-R.; Yang, C.-R.; Xu, M.; Zhang, Y.-J. Triterpenoids with Promoting Effects on

the Differentiation of PC12 Cells from the Steamed Roots of Panax notoginseng. *J. Nat. Prod*. **2015**, *78*(8), 1829-1840. https://doi.org/10.1021/acs.jnatprod.5b00027.

(8) Li, R.; Huang, T.; Nie, L.; Jia, A.; Zhang, L.; Yuan, Y.; Hong, Y.; Wang, J.; Hu, X. Chemical Constituents from Staminate Flowers of Eucommia ulmoides Oliver and Their Anti-Inflammation Activity in Vitro. *Chem. Biodivers*. **2021**, *18*(8), e2100331. https://doi.org/10.1002/cbdv.202100331.

(9) Wang, J.-R.; Yamasaki, Y.; Tanaka, T.; Kouno, I.; Jiang, Z.-H. Dammarane-type triterpene saponins from the flowers of Panax notoginseng. *Molecules*. **2009**, *14*(6), 2087-2094. https://doi.org/10.3390/molecules14062087.

(10) Xing, Q.; Liang, T.; Shen, G.; Wang, X.; Jin, Y.; Liang, X. Comprehensive HILIC × RPLC with mass spectrometry detection for the analysis of saponins in Panax notoginseng. *Analyst*. **2012**, *137*(9), 2239-2249. https://doi.org/10.1039/C2AN16078A.

(11) Yang, L.; Fang, Y.; Liu, R.; He, J. Phytochemical Analysis, Anti-inflammatory, and Antioxidant Activities of Dendropanax dentiger Roots. *Biomed. Res. Int*. **2020**, *2020*(22), 1-13. https://doi.org/10.1155/2020/5084057.

(12) Yang, Z.; Shao, Q.; Ge, Z.; Ai, N.; Zhao, X.; Fan X. A Bioactive Chemical Markers Based Strategy for Quality Assessment of Botanical Drugs: Xuesaitong Injection as a Case Study. *Sci. Rep*. **2017**, *7*(1), 2410. https://doi.org/10.1038/s41598-017-02305-y.

(13) Takemoto, T.; Arihara, S.; Yoshikawa, K. Studies on the Constituents of Cucurbitaceae Plants. XIV On the Saponin Constituents of Gynostemma pentaphyllum MAKINO.(9). Yakugaku Zasshi. **1986,** *104*(11), 1155-1162. https://doi.org/10.1248/yakushi1947.106.8_664.

(14) Cao, J.-L.; Ma, L.-J.; Wang, S.-P.; Deng, Y.; Wang, Y.-T.; Li, P.; Wan, J.B. Comprehensively qualitative and quantitative analysis of ginsenosides in Panax notoginseng leaves by online two-dimensional liquid chromatography coupled to hybrid linear ion trap Orbitrap mass spectrometry with deeply optimized dilution and modulation system. *Anal. Chim. Acta*. **2019**, *1079*, 237-251. https://doi.org/10.1016/j.aca.2019.06.040.

(15) Chen, J.; Tan, M.; Zou, L.; Liu, X.; Chen, S.; Shi, J.; Chen, C.; Wang, C.; Mei, Y.

Qualitative and Quantitative Analysis of the Saponins in Panacis Japonici Rhizoma Using Ultra-Fast Liquid Chromatography Coupled with Triple Quadrupole-Time of Flight Tandem Mass Spectrometry and Ultra-Fast Liquid Chromatography Coupled with Triple Quadrupole-Linear Ion Trap Tandem Mass Spectrometry. *Chem. Pharm. Bull (Tokyo)*. **2019**, *67*(8), 839-848. https://doi.org/10.1248/cpb.c19-00255.

(16) Shi, J.; Cai, Z.; Chen, S.; Zou, L.; Liu, X.; Tang, R.; Ma, J.; Wang, C.; Chen, J.; Tan, M. Qualitative and quantitative analysis of saponins in the flower bud of Panax ginseng (Ginseng Flos) by UFLC-Triple TOF-MS/MS and UFLC-QTRAP-MS/MS. *Phytochem. Anal*. **2020**, *31*(3), 287-296. https://doi.org/10.1002/pca.2894.

(17) Liu, X.-Y.; Wang, S.; Li, C.-J.; Ma, J.; Chen, F.-Y.; Peng, Y.; Wang, X.-L.; Zhang D.-M. Dammarane-type saponins from the leaves of Panax notoginseng and their neuroprotective effects on damaged SH-SY5Y cells. *Phytochemistry.* **2018**, *145*, 10-17. https://doi.org/10.1016/j.phytochem.2017.09.020.

(18) Zhao, J. Study on the cracking law and chemical characteristics of saponins in Panax notoginseng (Doctor Dissertation). *Beijing University of Chinese Medicine*. **2015**, 201. https://kns.cnki.net/KCMS/detail/detail.aspx?dbname=CMFD201502&filename=101 5390183.nh.

(19) Zuo, T.; Qian, Y.; Zhang, C.; Wei, Y.; Wang, X.; Wang, H.; Hu, Y.; Li, W.; Wu, X.; Yang, W. Data-Dependent Acquisition and Database-Driven Efficient Peak Annotation for the Comprehensive Profiling and Characterization of the Multicomponents from Compound Xueshuantong Capsule by UHPLC/IM-QTOF-MS. *Molecules*. **2019**, *24*(19), 3431. https://doi.org/10.3390/molecules24193431.

(20) Liu, J.-H.; Wang, X.; Cai, S. Q.; Komatsu, K.; Namba, T. Analysis of the Constituents in the Chinese Drug Notoginseng by Liquid Chromatography-Electrospray Mass Spectrometry. *J. Chin. Pharm. Sci*. **2004**, *13*(4), 225-237. https://kns.cnki.net/kcms/detail/detail.aspx?FileName=XYGZ200404000&DbName=CJFQ2004.

(21) Zhou，Q.-L.; Xu, W.; Yang, X.W. Chemical constituents of Chinese red ginseng. *Zhongguo Zhong Yao Za Zhi*. **2016**, *41*(2), 233-249.

https://schlr.cnki.net/en/Detail/index/GARJ2016/SJPD5530B7DC7CEB1652B6DE33AAEB27FA8C.
